# Acute social defeat induces cannabinoid type 1 receptor-dependent synaptic depression in the anterior ventral subiculum to anterior hypothalamic nucleus pathway to drive social avoidance in male mice

**DOI:** 10.64898/2026.10.25.760350

**Authors:** Changsu Woo, Soonje Lee, Changwoo Lee, Donghyeon Yoon, Sukwoo Choi, Shin Jung Kang, Kyuhyun Choi, Ki Soon Shin

## Abstract

**BACKGROUND:** Social avoidance is a protective response typically studied after repeated defeat. A single episode of acute social defeat stress (ASDS) is sufficient to trigger this avoidance. However, the specific circuit modifications underlying this rapid behavioral shift remain unknown. A candidate for this rapid regulation is the pathway from the ventral subiculum (vSUB), a major hippocampal output, to the anterior hypothalamic nucleus (AHN), a critical hub for stress responses.

**METHODS:** Male mice underwent ASDS or subthreshold social defeat stress. We combined social interaction, open field, and elevated plus maze testing with optogenetic-assisted slice recording, pharmacology, pathway-specific chemogenetics and optogenetics, retrograde and anterograde transsynaptic tracing, and in situ hybridization.

**RESULTS:** ASDS induced presynaptic depression at anterior vSUB (avSUB)→AHN synapses through a glucocorticoid-and cannabinoid type 1 receptor (CB1R)-dependent mechanism, while posterior vSUB (pvSUB)→AHN synapses were unaffected. Infusing a CB1R antagonist into the AHN before ASDS attenuated both the synaptic depression and the social avoidance. Chemogenetic activation of the avSUB→AHN pathway during ASDS attenuated social avoidance, whereas its inhibition induced avoidance under subthreshold stress. Optogenetic activation of the avSUB→AHN pathway produced an immediate increase in social approach. Anterograde tracing revealed that anterior and posterior vSUB neurons project to largely separate AHN populations. The pvSUB→AHN pathway modulates anxiety-like behavior without altering social approach.

**CONCLUSIONS:** These findings reveal specialized roles for vSUB subregions in social and emotional responses, and identify presynaptic gating of the avSUB→AHN synapse as a control point for stress-induced social withdrawal.

## INTRODUCTION

Social avoidance is a spectrum of behaviors through which an individual withdraws from or evades social interaction, occurs across species, and is provoked by aversive social experiences such as rejection, defeat, or conflict (1). In moderation it is adaptive, because it limits exposure to threatening social stimuli. When it becomes persistent and generalized it promotes isolation and is a prominent feature of depression, anxiety disorders, and post-traumatic stress disorder (1). Social defeat reliably elicits social avoidance and has become the leading model for studying its neural basis (2–4). Most mechanistic work has examined adaptations accumulating over repeated defeat, yet a single episode, acute social defeat stress (ASDS), is sufficient to produce avoidance (5). What remains unclear is which defined circuit is modified by a single acute defeat, and whether that change is required for the avoidance that follows.

A strong candidate is the projection from the ventral subiculum (vSUB) to the anterior hypothalamic nucleus (AHN). The vSUB is the principal output station of the hippocampal formation and a key node in the stress response (6,7), and acute stress rapidly modifies transmission at vSUB and ventral hippocampal synapses (8,9). The AHN is a component of the medial hypothalamic defensive network that converts threat-related information into coordinated output (10–12) and contributes to regulation of the stress axis (13). It is predominantly GABAergic (14), hippocampal afferents supply a large fraction of the extrahypothalamic input to its GABAergic neurons (13), and vSUB neurons constitute the principal hippocampal excitatory drive onto that population (15). The vSUB→AHN pathway is therefore positioned to translate a stressful social experience into defensive behavioral output.

The behavioral role established so far for the vSUB→AHN pathway is not a social one. A recent study showed that silencing vSUB input to AHN GABAergic neurons reduces avoidance, but the avoidance examined was innate and nonsocial, evoked by objects and predator cues (15). The AHN GABAergic population also did not change its activity while mice investigated a conspecific (15). The stereotaxic coordinates in that study place the injections in the posterior vSUB (pvSUB), leaving the possibility that a distinct pathway arising from the anterior vSUB (avSUB) carries social information.

Subicular projections to the hypothalamus are topographically organized along the longitudinal axis (16), and vSUB projection neurons comprise transcriptomically distinct classes with dissociable outputs (17,18). Different portions of the vSUB may therefore reach one hypothalamic target through separate channels. Functionally, social avoidance and general anxiety are separable states that can be dissociated by circuit manipulation (19,20), and vSUB output can increase or decrease avoidance depending on its downstream target (21).

A single stressful experience is sufficient to reset excitatory transmission at defined synapses (22–24). In stress-responsive regions that change is frequently expressed presynaptically rather than as a postsynaptic adjustment (23,24). This rapid, presynaptically expressed depression is implemented by endocannabinoids (eCB), which are released postsynaptically and act retrogradely on presynaptic cannabinoid type 1 receptors (CB1R) to lower release probability (25–29). In the paraventricular nucleus of the hypothalamus (PVN), this pathway is engaged by glucocorticoids, which mobilize eCB through a nongenomic mechanism and suppress glutamate release within minutes (30–32), and acute stress recruits the same signaling in the amygdala (33,34). Because brain corticosterone peaks within the first half hour of a stressor, later than in plasma (35), this mechanism is positioned to act while the stressor is still in progress. The eCB system also regulates the stress response and social behavior (36,37) and disrupting CB1R signaling alters social avoidance and anxiety-like responses toward conspecifics (38,39).

Here, we show that a single episode of ASDS selectively reduces glutamate release at avSUB→AHN but not pvSUB→AHN synapses, that corticosterone reproduces this depression, and that local blockade of CB1R in the AHN attenuates both the depression and the social avoidance. Chemogenetic manipulation of this pathway bidirectionally alters social behavior, and optogenetic activation acutely increases social approach. We further identify a topographic dissociation in which the avSUB→AHN pathway preferentially targets non-GABAergic AHN neurons and regulates social approach, whereas the pvSUB→AHN pathway modulates anxiety-like behavior without altering social approach.

## METHODS AND MATERIALS

Full protocols, stereotaxic coordinates, viral vectors, drug doses, recording parameters, histological procedures and the criteria used to select each statistical test are provided in the Supplement.

### Animals

All experiments were approved by the Kyung Hee University Animal Care and Use Committee (KHSASP-23-407, KHSASP-24-584, and KHSASP-25-680). Adult male wild-type C57BL/6N mice and male *Gad2*-IRES-Cre mice were used, and male CD-1 retired breeder mice served as aggressors. Only male mice were studied.

### Stress Protocols and Behavioral Testing

For ASDS, the test mouse was physically defeated by a screened CD-1 aggressor for 10 min and then kept in the same cage behind a perforated Plexiglas partition for a further 20 min of sensory exposure, giving a 30-min protocol. Subthreshold social defeat stress (SSDS) consisted of 5 min of physical defeat followed by 25 min in the animal’s own home cage, and lacked the sensory exposure phase. Social interaction was tested in an open field containing a target cage, first without and then with an unfamiliar CD-1 target, and anxiety-like behavior was assessed in the open field test (OFT) and the elevated plus maze (EPM).

### Pathway-Specific Manipulation and Electrophysiology

Viral vectors were injected into the avSUB, the pvSUB, or the AHN, and guide cannulae or optic fibers were implanted over the AHN. To isolate the subicular input to the AHN, the opsin Chronos was expressed in the avSUB or the pvSUB and light-evoked excitatory postsynaptic currents were recorded from AHN neurons in acute slices. Presynaptic release was assessed from the paired-pulse ratio (PPR).

### Anterograde Transsynaptic Tracing and Histology

To determine whether the avSUB and the pvSUB innervate the same AHN neurons, AAV1 carrying Cre was injected into one subregion and AAV1 carrying FlpO into the other, together with Cre-dependent and Flp-dependent reporters in the AHN, and single-and double-labeled neurons were counted. The postsynaptic cell type of the anterior pathway was examined in *Gad2*-IRES-Cre mice and by RNAscope in situ hybridization for *Slc17a6* (*Vglut2*) and *Slc32a1* (*Vgat*).

### Statistics

Data are mean ± SEM and significance was set at p <.05. Two-group comparisons used t tests, comparisons among three or more groups used one-way analysis of variance (ANOVA), and designs in which the same animal was tested under more than one condition used two-way repeated-measures ANOVA, each followed by the appropriate post hoc test. Analyses were performed with GraphPad Prism 10.0. The exact n, test statistics and p values for every comparison are reported in Supplemental Table S1.

## RESULTS

### ASDS INDUCES SOCIAL AVOIDANCE AND THE avSUB→AHN PATHWAY SUPPORTS SOCIAL APPROACH

Male C57BL/6N mice underwent a single session of ASDS and were tested in the social interaction test (SIT) immediately afterwards (Figure 1A). Naive mice spent more time in the interaction zone when the target was introduced, whereas defeated mice did not, and interaction time in the presence of the target was significantly lower in defeated mice (Figure 1B, Figure S1A). Interaction time without a target did not differ between groups, so the effect was specific to the social stimulus (Figure 1B). Corner occupancy changed in opposite directions in the two groups (Figure 1C). Social interaction ratios and social preference indices were also lower in defeated mice (Figure S1B, S1C). Locomotion was reduced after ASDS, particularly with the target present (Figure 1D). The group difference persisted with distance moved as a covariate (Figure S1D), so reduced locomotion does not account for the reduced social interaction. Social avoidance was still present when the same mice were retested 24 hours later (Figure 1E).

**Figure 1.**
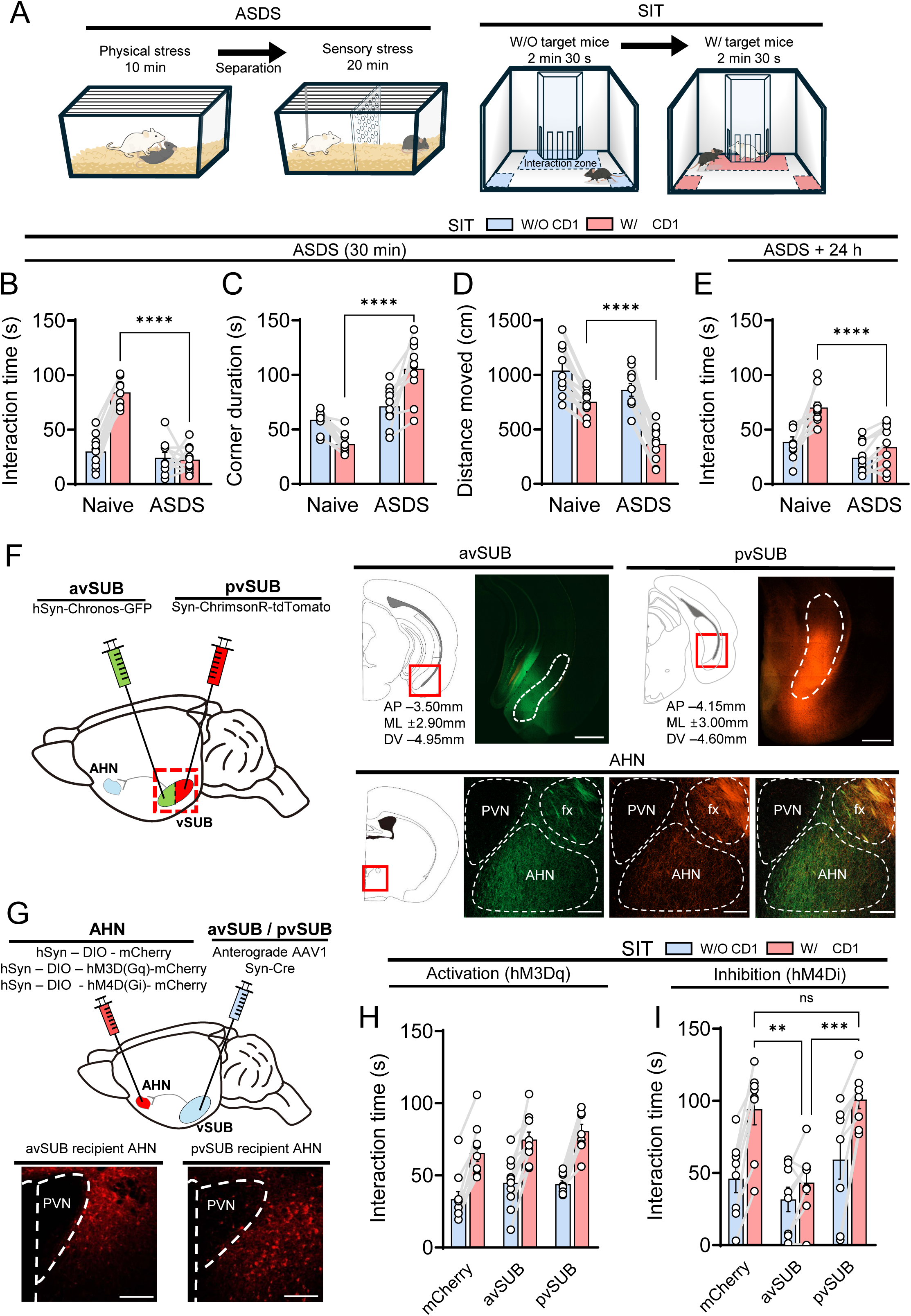
Acute social defeat stress (ASDS) induces social avoidance, and the anterior ventral subiculum (avSUB)→anterior hypothalamic nucleus (AHN) pathway supports social approach. (A) Schematic of the ASDS protocol and of the social interaction test (SIT). Mice received 10 min of physical interaction with an aggressive CD-1 mouse followed by 20 min of sensory exposure across a perforated divider, and were tested immediately afterwards in two consecutive 2 min 30 s epochs, first without and then with a novel CD-1 target. (B) Time spent in the interaction zone (naive versus ASDS with the target present, p <.0001, n = 10 and 10 mice). (C) Time spent in the corners (naive versus ASDS with the target present, p <.0001, n = 10 and 10 mice). (D) Distance moved (naive versus ASDS with the target present, p <.0001, group by target interaction p =.025, n = 10 and 10 mice). (E) Time spent in the interaction zone 24 hours after ASDS (naive versus ASDS with the target present, p <.0001, group by target interaction p =.026, n = 10 and 10 mice). (F) Dual anterograde labeling of the anterior and posterior ventral subiculum (vSUB). AAV-hSyn-Chronos-GFP was injected into the avSUB and AAV-Syn-ChrimsonR-tdTomato into the posterior vSUB (pvSUB) of the same animals. The representative images show the two injection sites with their stereotaxic coordinates and the resulting terminal fields in the AHN, in which the paraventricular nucleus of the hypothalamus (PVN) and the fornix (fx) are shown as anatomical landmarks. (G) Strategy for chemogenetic control of AHN neurons receiving avSUB or pvSUB input. Anterograde transsynaptic AAV1-hSyn-Cre was injected into either the avSUB or the pvSUB and Cre-dependent hM3D(Gq), hM4D(Gi), or mCherry into the AHN. The representative images show mCherry-labeled AHN neurons after avSUB or pvSUB injection, with the PVN shown as an anatomical landmark. (H) Interaction time in stress-naive mice after chemogenetic activation with hM3D(Gq) (no effect of group and no group by target interaction, n = 10 mCherry, 10 avSUB and 9 pvSUB mice). (I) Interaction time in stress-naive mice after chemogenetic inhibition with hM4D(Gi) (with the target present, mCherry versus avSUB p =.002 and avSUB versus pvSUB p <.001, n = 8 mice per group). Light blue bars show the epoch without a target (W/O) and light red bars the epoch with a target (W/). Open circles are individual mice and gray lines connect the two epochs within a mouse. Data are mean ± SEM. Panels B to E were analyzed by two-way repeated measures ANOVA with Šídák’s multiple comparisons test, and panels H and I by two-way repeated measures ANOVA with Tukey’s multiple comparisons test. Statistics are reported in Table S1. In all figures, *p <.05, **p <.01, ***p <.001 and ****p <.0001, and ns denotes not significant. Scale bars, 500 μm in the subicular images of (F) and 200 μm in all other images.

We found that the vSUB sends dual projections to the AHN originating from its anterior and posterior subregions (Figure 1F). We thus asked whether these two pathways (avSUB→AHN and pvSUB→AHN) contribute differently to social approach. To selectively target AHN neurons innervated by each population, we injected anterograde transsynaptic AAV1-hSyn-Cre into either the avSUB or the pvSUB, and delivered Cre-dependent designer receptors exclusively activated by designer drugs (DREADDs, hM3D(Gq) or hM4D(Gi)) or mCherry (as a control) in the AHN (40–42) (Figure 1G). This strategy favors release-dependent Cre transfer to postsynaptic AHN neurons. In stress-naive mice, chemogenetic activation of either postsynaptic population did not alter baseline social approach (Figure 1H, Figure S5A). Meanwhile, inhibition of AHN neurons receiving avSUB input reduced interaction time (Figure 1I) and distance moved (Figure S5B) in the presence of a target, whereas inhibition of AHN neurons receiving pvSUB input had no effect (Figure 1I, Figure S5B). These results reveal that silencing the avSUB→AHN pathway is sufficient to reduce social approach in the absence of any stressor.

### ASDS SELECTIVELY REDUCES RELEASE PROBABILITY AT avSUB→AHN, BUT NOT pvSUB→AHN, SYNAPSES

To determine whether ASDS alters synaptic transmission of the vSUB→AHN pathways, we expressed the highly sensitive channelrhodopsin variant Chronos (43) in the avSUB (Figure 2A) or pvSUB (Figure 2D) and recorded light-evoked currents from AHN neurons ex vivo. Synaptic transmission across both avSUB→AHN and pvSUB→AHN pathways was completely blocked by glutamate receptor antagonists, D-APV and CNQX, indicating excitatory glutamatergic transmission (Figure 2A, 2D). In slices from defeated mice, light-evoked current amplitude in the avSUB→AHN pathway was reduced relative to naive mice (Figure 2B) and the paired-pulse ratio (PPR) was increased (Figure 2C), indicating a presynaptic depression of glutamate release after ASDS (44). Systemic administration of the glucocorticoid receptor antagonist mifepristone (RU486) before defeat prevented both ASDS-induced changes (Figure 2B, 2C), demonstrating that glucocorticoid signaling is required for the effect. A time-course analysis of slices prepared at 30 min, 90 min, 3 hours and 24 hours post-ASDS revealed that both light-evoked current amplitude and the PPR at avSUB→AHN synapses progressively returned to control levels, reaching full recovery by 24 hours (Figure S2). When the same measurements were made in the pvSUB→AHN pathway, however, light-evoked current amplitude did not differ between naive and defeated mice (Figure 2E) and the PPR remained unaltered (Figure 2F). Together, these results demonstrate that ASDS selectively reduces release probability at avSUB→AHN synapses while sparing the pvSUB→AHN pathway. These data suggest that while the avSUB→AHN synaptic depression is readily reversible, it may act as an acute trigger to initiate persistent behavioral changes rather than maintaining them directly (Figure 1B, 1E).

**Figure 2.**
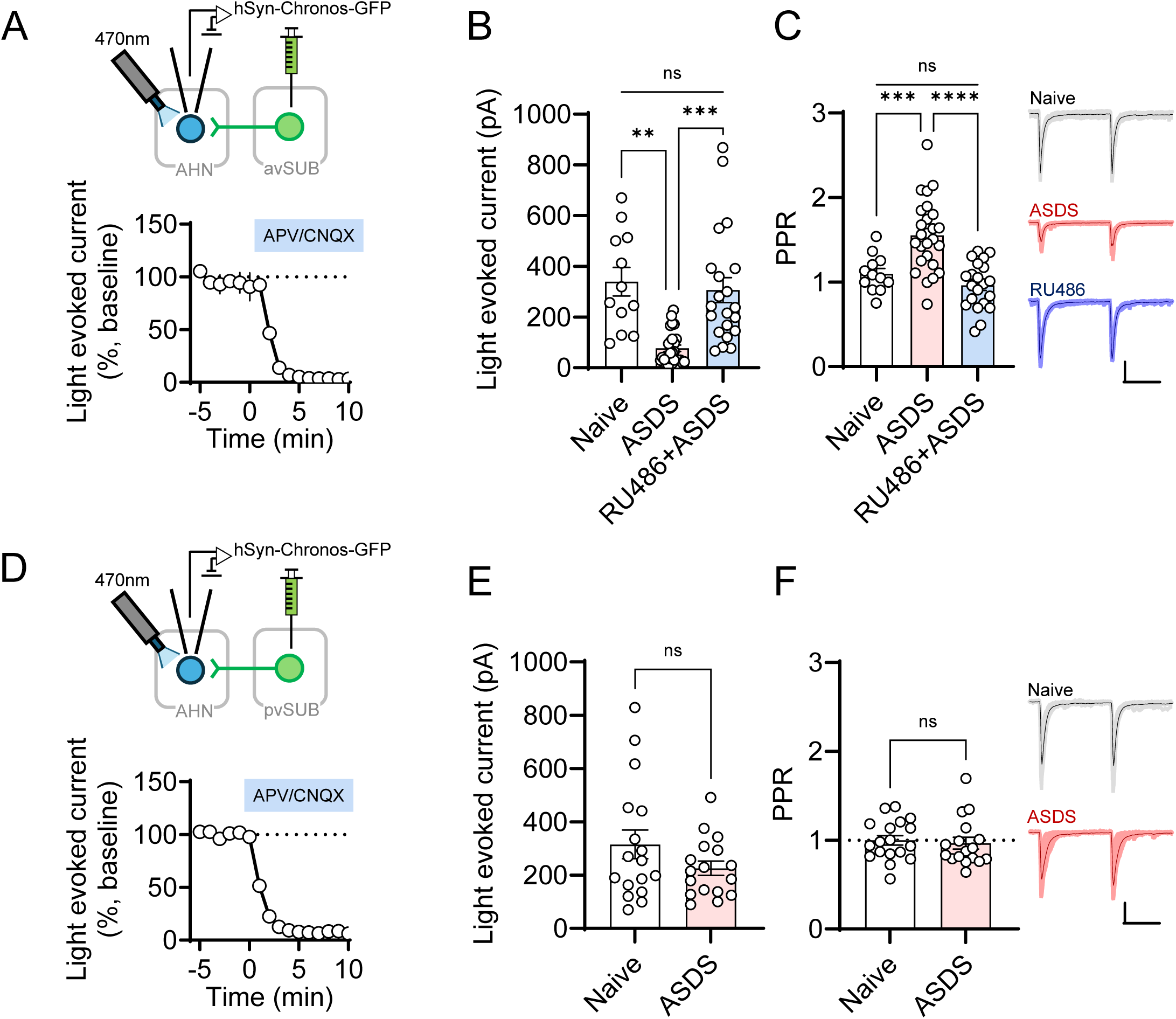
Acute social defeat stress (ASDS) selectively reduces release probability at anterior ventral subiculum (avSUB)→anterior hypothalamic nucleus (AHN) synapses. (A) Strategy for recording the avSUB→AHN pathway. AAV-hSyn-Chronos-GFP was injected into the avSUB and light-evoked currents were recorded from AHN neurons in acute slices. The lower panel shows the block of the light-evoked current by D-2-amino-5-phosphonovaleric acid (D-APV) and 6-cyano-7-nitroquinoxaline-2,3-dione (CNQX). (B) Light-evoked current amplitude in naive mice, mice exposed to ASDS, and mice given mifepristone (RU486) before ASDS (naive versus ASDS p =.002, ASDS versus RU486+ASDS p <.001, naive versus RU486+ASDS not significant, n = 12, 26 and 21 cells from 5, 7 and 4 mice). (C) Paired-pulse ratio (naive versus ASDS p <.001, ASDS versus RU486+ASDS p <.0001, naive versus RU486+ASDS not significant, n as in B). Representative traces are shown at right. (D) Strategy for recording the posterior ventral subiculum (pvSUB)→AHN pathway, as in (A). (E) Light-evoked current amplitude (naive versus ASDS p =.150, n = 17 and 17 cells from 3 and 4 mice). (F) Paired-pulse ratio (naive versus ASDS p =.699, n as in E). Representative traces are shown at right. Data are mean ± SEM. Panels B and C were analyzed by Brown-Forsythe and Welch ANOVA with Dunnett’s T3 multiple comparisons test, and panels E and F by unpaired t test with Welch’s correction. Statistics are reported in Table S1. Scale bars, 100 pA and 50 ms.

### GLUCOCORTICOID SUPPRESSES GLUTAMATERGIC TRANSMISSION IN THE avSUB→AHN PATHWAY VIA CB1R

Glucocorticoids are well known to suppress presynaptic glutamate release in the hypothalamus by mobilizing eCB (30–32). Therefore, we asked whether ex vivo corticosterone application is sufficient to induce synaptic depression in the avSUB→AHN pathway and whether this process is eCB-dependent. Slices from naive mice expressing Chronos in the avSUB were incubated with corticosterone, which reduced light-evoked current amplitude and increased the PPR, recapitulating the changes observed after ASDS (Figure 3A, 3B). Coapplication of the CB1R antagonist AM251 with corticosterone prevented both effects (Figure 3A, 3B). We also found that application of the CB1R agonist ACEA reduced light-evoked current amplitude (Figure 3C) and increased the PPR (Figure 3D). These findings indicate not only that CB1R mediates corticosterone-induced synaptic depression, but also that direct activation of CB1R is sufficient to drive this presynaptic depression.

**Figure 3.**
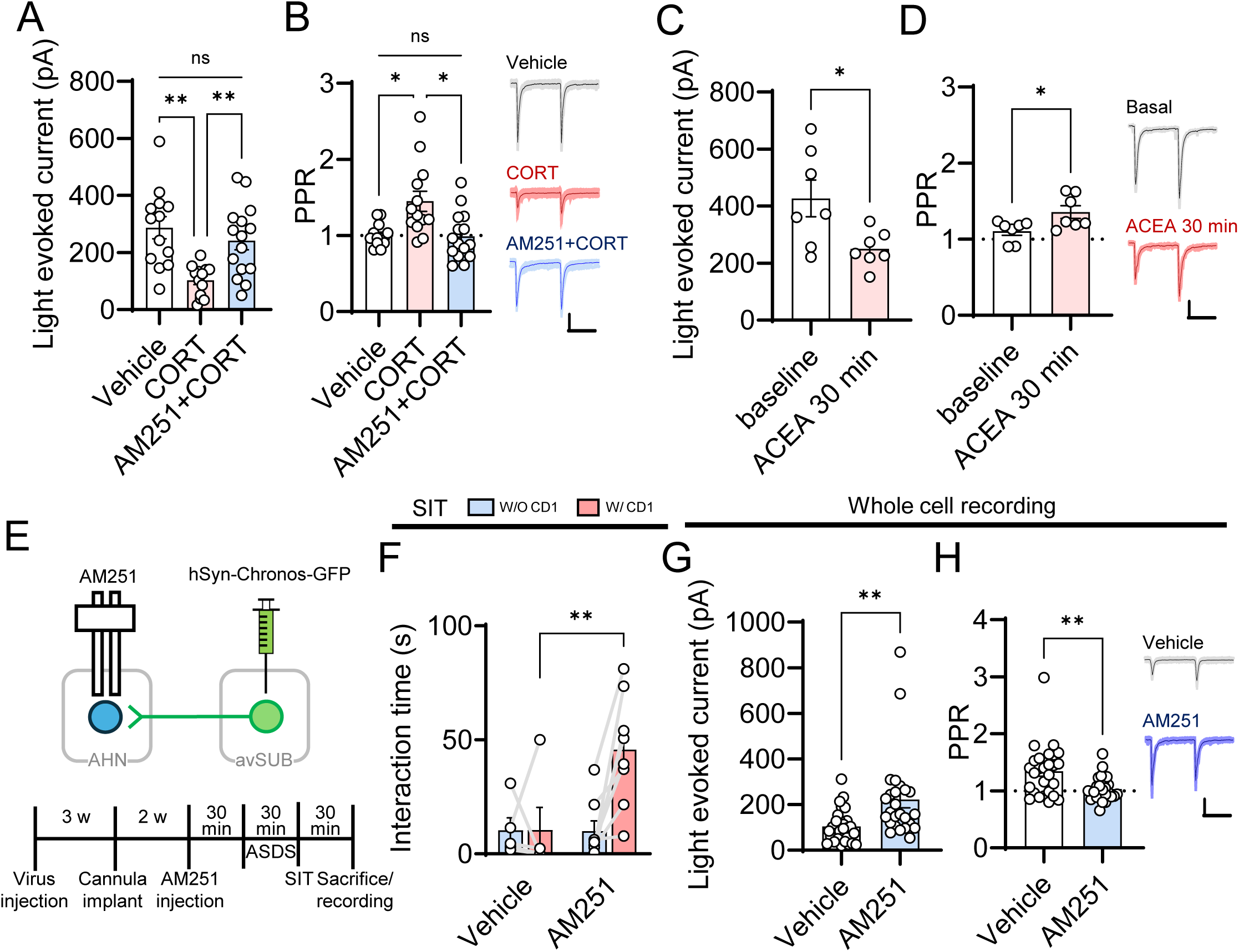
Corticosterone suppresses glutamate release at anterior ventral subiculum (avSUB)→anterior hypothalamic nucleus (AHN) synapses through cannabinoid type 1 receptor (CB1R). (A) Light-evoked current amplitude in slices from naive mice after bath application of vehicle, corticosterone, or corticosterone together with AM251 (vehicle versus corticosterone p =.001, corticosterone versus AM251+corticosterone p =.003, vehicle versus AM251+corticosterone not significant, n = 13, 13 and 15 cells from 5, 4 and 4 mice). (B) Paired-pulse ratio in the same recordings (vehicle versus corticosterone p =.019, corticosterone versus AM251+corticosterone p =.023, vehicle versus AM251+corticosterone not significant, n as in A). Representative traces are shown at right. (C) Light-evoked current amplitude before and 30 min after application of the CB1R agonist ACEA (p =.028, n = 7 cells from 4 mice). (D) Paired-pulse ratio in the same recordings (p =.015, n as in C). Representative traces are shown at right. (E) Timeline and strategy for local AM251 infusion into the AHN. A cannula was implanted above the AHN and AM251 or vehicle was infused before acute social defeat stress (ASDS), after which mice were tested in the social interaction test (SIT) and slices were prepared from the same animals. (F) Interaction time after ASDS (vehicle versus AM251 with the target present p =.007, n = 5 and 8 mice). (G) Light-evoked current amplitude in slices from the mice in (F) (vehicle versus AM251 p =.006, n = 27 and 25 cells from 4 mice per group). (H) Paired-pulse ratio in the same recordings (vehicle versus AM251 p =.003, n as in G). Representative traces are shown at right. Data are mean ± SEM. Panels A and B were analyzed by Brown-Forsythe and Welch ANOVA with Dunnett’s T3 multiple comparisons test, panels C and D by paired t test, panel F by two-way repeated measures ANOVA with Šídák’s multiple comparisons test, and panels G and H by unpaired t test with Welch’s correction. Statistics are reported in Table S1. Scale bars, 100 pA and 50 ms.

To test whether this eCB-CB1R signaling underlies the ASDS-induced synaptic depression and social avoidance in vivo, we locally infused AM251 into the AHN prior to ASDS, assessed the mice in the SIT, and subsequently prepared acute slices from the same animals for electrophysiological recordings (Figure 3E). Local AM251 infusion attenuated the ASDS-induced social avoidance, with AM251-infused mice spending more time in the interaction zone with the target present than vehicle controls (Figure 3F), and distance moved did not differ (Figure S5C). In acute slices prepared from these same mice, light-evoked current amplitudes were larger and the PPR was lower in the AM251 group compared to vehicle-treated controls (Figure 3G, 3H). Thus, blocking CB1R within the AHN attenuates both the synaptic depression and the social avoidance, confirming that corticosterone-driven retrograde eCB signaling is the critical mediator of mediates the ASDS response.

### CHEMOGENETIC MANIPULATION OF THE avSUB→AHN PATHWAY DURING DEFEAT PRODUCES LONG-LASTING EFFECTS ON SOCIAL AVOIDANCE

We next manipulated the activity of avSUB neurons projecting to the AHN. To selectively target this neuronal population, we injected AAVretro-EF1a-Cre (45) into the AHN and Cre-dependent DREADDs (hM3D(Gq) or hM4D(Gi)) into the avSUB (Figure 4A). We administered clozapine N-oxide (CNO) or vehicle 30 min prior to the stressor. Mice underwent the SIT immediately after ASDS and were re-tested 24 hours later without further ASDS. Chemogenetic activation of these neurons attenuated ASDS-induced social avoidance, and this protective effect persisted 24 hours later (Figure 4B, 4C). Because the standard ASDS protocol suppresses social interaction to near-floor levels, any further decrease driven by chemogenetic manipulation would be difficult to detect. Therefore, we employed subthreshold social defeat stress (SSDS). This protocol incorporated the physical and social stress of an aggressor encounter but failed to induce social avoidance on its own (Figure S3). Remarkably, inhibition of AHN-projecting avSUB neurons during SSDS was sufficient to produce social avoidance, an effect that persisted 24 hours later (Figure 4D, 4E).

**Figure 4.**
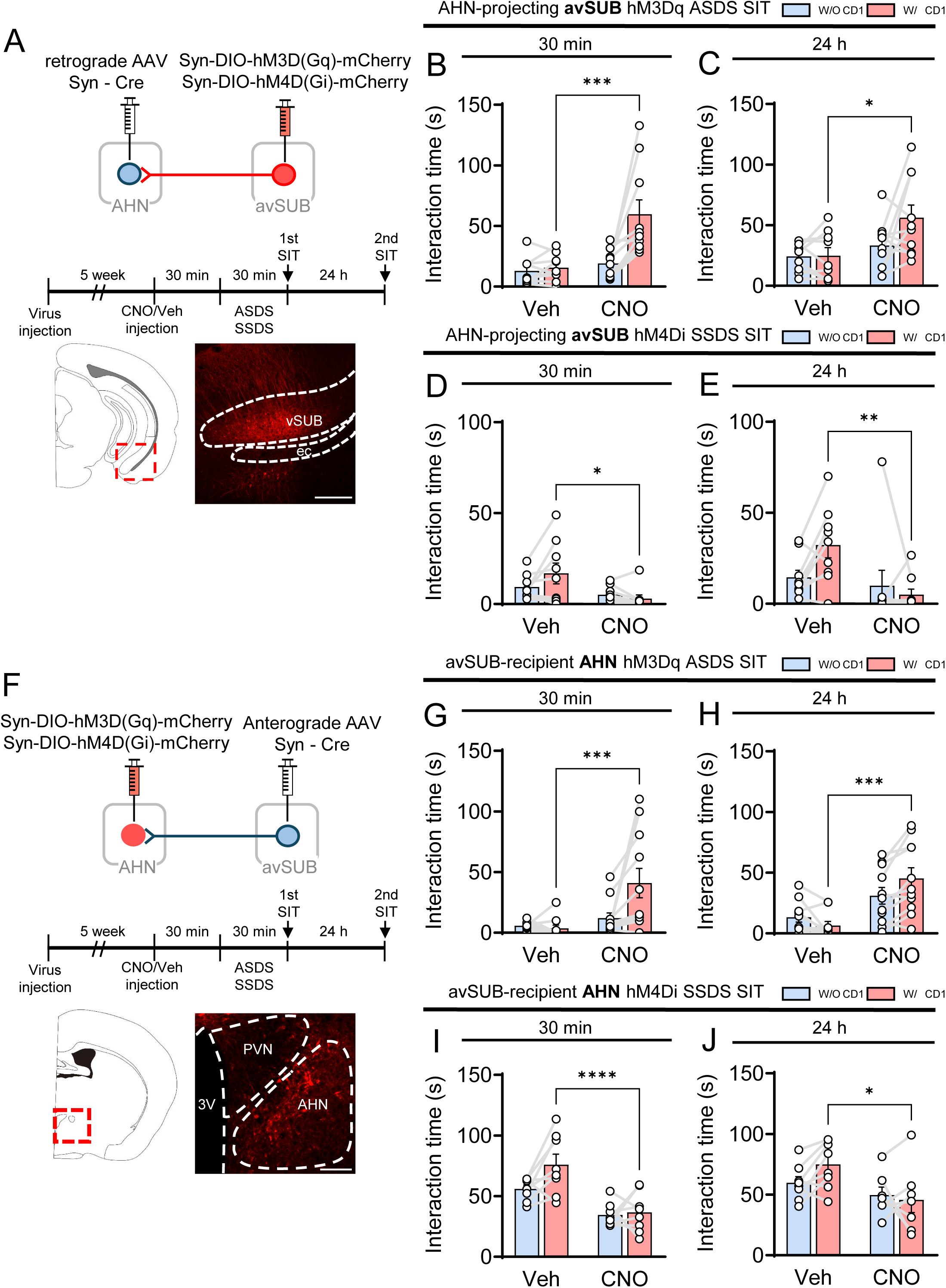
Chemogenetic manipulation of the anterior ventral subiculum (avSUB)→anterior hypothalamic nucleus (AHN) pathway during defeat produces long-lasting changes in social behavior. (A) Strategy and timeline for chemogenetic control of AHN-projecting avSUB neurons. AAVretro-EF1a-Cre was injected into the AHN and Cre-dependent hM3D(Gq), hM4D(Gi), or mCherry into the avSUB. Clozapine N-oxide (CNO) or vehicle was given 30 min before the stressor, and mice were tested in the social interaction test (SIT) immediately afterwards and again 24 hours later. The representative image shows mCherry-labeled avSUB neurons. (B) Interaction time in the first test after acute social defeat stress (ASDS) with hM3D(Gq) (vehicle versus CNO with the target present p <.001, n = 9 and 10 mice). (C) Interaction time in the second test 24 hours later (p =.011, n = 9 and 10 mice). (D) Interaction time in the first test after subthreshold social defeat stress (SSDS) with hM4D(Gi) (p =.012, n = 9 and 9 mice). (E) Interaction time in the second test 24 hours later (p =.006, n = 9 and 9 mice). (F) Strategy and timeline for chemogenetic control of AHN neurons receiving avSUB input. Anterograde transsynaptic AAV1-hSyn-Cre was injected into the avSUB and Cre-dependent hM3D(Gq), hM4D(Gi), or mCherry into the AHN. The representative image shows mCherry-labeled AHN neurons. (G) Interaction time in the first test after ASDS with hM3D(Gq) (p <.001, n = 10 and 11 mice). (H) Interaction time in the second test 24 hours later (p <.001, n = 10 and 11 mice). (I) Interaction time in the first test after SSDS with hM4D(Gi) (p <.0001 with the target present, n = 8 and 8 mice). (J) Interaction time in the second test 24 hours later (p =.020, n = 8 and 7 mice). Light blue bars show the epoch without a target and light red bars the epoch with a target. Open circles are individual mice and gray lines connect the two epochs within a mouse. Data are mean ± SEM. All panels were analyzed by two-way repeated measures ANOVA with Šídák’s multiple comparisons test. Statistics are reported in Table S1. Scale bars, 200 μm.

Manipulating the activity of the postsynaptic AHN neurons that receive avSUB input yielded comparable behavioral outcomes. To selectively target this neuronal population, we injected anterograde AAV1-hSyn-Cre into the avSUB and Cre-dependent DREADDs into the AHN (Figure 4F). Chemogenetic activation of AHN neurons receiving avSUB input during ASDS attenuated social avoidance, which persisted 24 hours later (Figure 4G, 4H). Although SSDS alone failed to induce social avoidance, inhibition during SSDS produced social avoidance that persisted 24 hours later (Figure 4I, 4J). It is noteworthy that a single administration of CNO prior to defeat altered social interaction not only immediately afterwards but also 24 hours later, a time point at which CNO is no longer pharmacologically active (46). Data for distance moved across all aforementioned SIT are presented in Figure S5. CNO alone did not alter social interaction in mice lacking DREADD expression (Figure S4). Overall, our results indicate that the level of avSUB→AHN pathway activity during stress exposure determines the development of persistent social avoidance.

### OPTOGENETIC ACTIVATION OF THE avSUB→AHN PATHWAY ATTENUATES ASDS-INDUCED SOCIAL AVOIDANCE

Because the slow clearance of CNO precludes time-locked measurements, we next used optogenetic activation of the avSUB→AHN pathway. Retrograde AAV-CAG-Cre was injected into the AHN together with Cre-dependent AAV-hSyn-FLEX-Chronos-GFP into the avSUB, which restricted Chronos expression to AHN-projecting avSUB neurons, and an optical fiber was implanted above the AHN (Figure 5A, 5B). The SIT comprised five epochs in which the light alternated between off and on (Figure 5B). In stress-naive animals, social interaction time further increased during light-on stimulation in Chronos-expressing mice, whereas interaction time in GFP-expressing controls steadily declined over the course of the test (Figure 5C).

**Figure 5.**
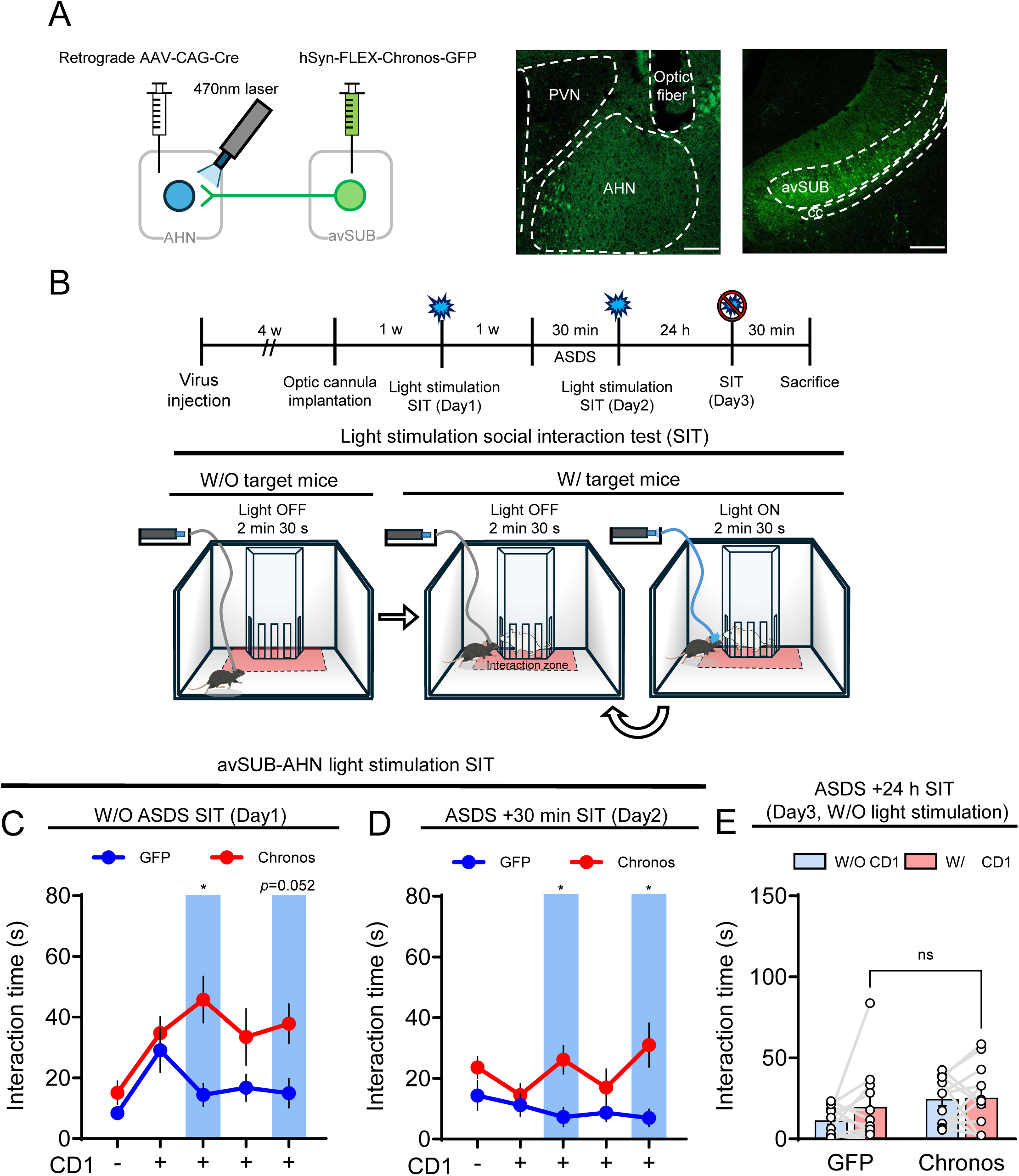
Optogenetic activation of the anterior ventral subiculum (avSUB)→anterior hypothalamic nucleus (AHN) pathway produces a time-locked attenuation of social avoidance. (A) Strategy for optogenetic stimulation of avSUB terminals in the AHN. Retrograde AAV-CAG-Cre was injected into the AHN together with Cre-dependent AAV-hSyn-FLEX-Chronos-GFP into the avSUB, and an optical fiber was implanted above the AHN. The representative images show Chronos-GFP expression in the avSUB and labeled terminals together with the optic fiber tract in the AHN. (B) Timeline of the experiment and design of the social interaction test (SIT), which comprised five consecutive 2 min 30 s epochs, a baseline epoch without a target and with the light off followed by four epochs with the target present in which the light alternated between off and on. (C) Interaction time across the five epochs in stress-naive mice (GFP versus Chronos p =.010 in the first light-on epoch and p =.052 in the second, n = 11 and 12 mice). (D) Interaction time across the five epochs 30 min after acute social defeat stress (ASDS) (GFP versus Chronos p =.016 in the first light-on epoch and p =.038 in the second, n = 11 and 12 mice). (E) Interaction time in the test performed 24 hours after ASDS without optogenetic stimulation (GFP versus Chronos with the target present p =.688, n = 12 and 12 mice). Blue shading marks the light-on epochs. Data are mean ± SEM. Panels C to E were analyzed by two-way repeated measures ANOVA with Šídák’s multiple comparisons test, with the Geisser-Greenhouse correction applied in panels C and D. Statistics are reported in Table S1. Scale bars, 200 μm.

We next asked whether acute optogenetic activation of the avSUB→AHN pathway could reverse social avoidance established by ASDS. Indeed, optogenetic stimulation significantly attenuated social avoidance in Chronos-expressing mice, driving an increase in interaction time during the light-on epochs (Figure 5D). This rescue was strictly dependent on the stimulation, because interaction time declined to the level of GFP-expressing controls whenever the light was off. When re-tested 24 hours later without optogenetic stimulation, both Chronos-and GFP-expressing mice avoided the social target to a similar extent (Figure 5E). Distance moved did not differ between Chronos-and GFP-expressing mice in the test performed 24 hours after ASDS (Figure S5D).

### avSUB AND pvSUB PREFERENTIALLY TARGET SEPARATE POPULATIONS OF AHN NEURONS

Having established that ASDS selectively depresses the avSUB→AHN pathway to drive social avoidance, leaving the pvSUB→AHN pathway unaffected, we next investigated the underlying anatomical differences between these two pathways. We injected transsynaptic anterograde AAV1-EF1a-FlpO into the avSUB and anterograde AAV1-hSyn-Cre into the pvSUB, respectively. Concurrently, we delivered Flp-dependent mCherry and Cre-dependent EYFP into the AHN, labeling the postsynaptic targets of both subregions in the same brain (47) (Figure 6A, 6B). Among all labeled AHN neurons, 27.4% received avSUB input alone, 48.5% received pvSUB input alone, and 24.1% received both (Figure 6C). Thus, most postsynaptic AHN neurons are innervated by only one of these pathways, demonstrating that the avSUB and pvSUB target largely separate neuronal populations within the AHN (Figure 6D).

**Figure 6.**
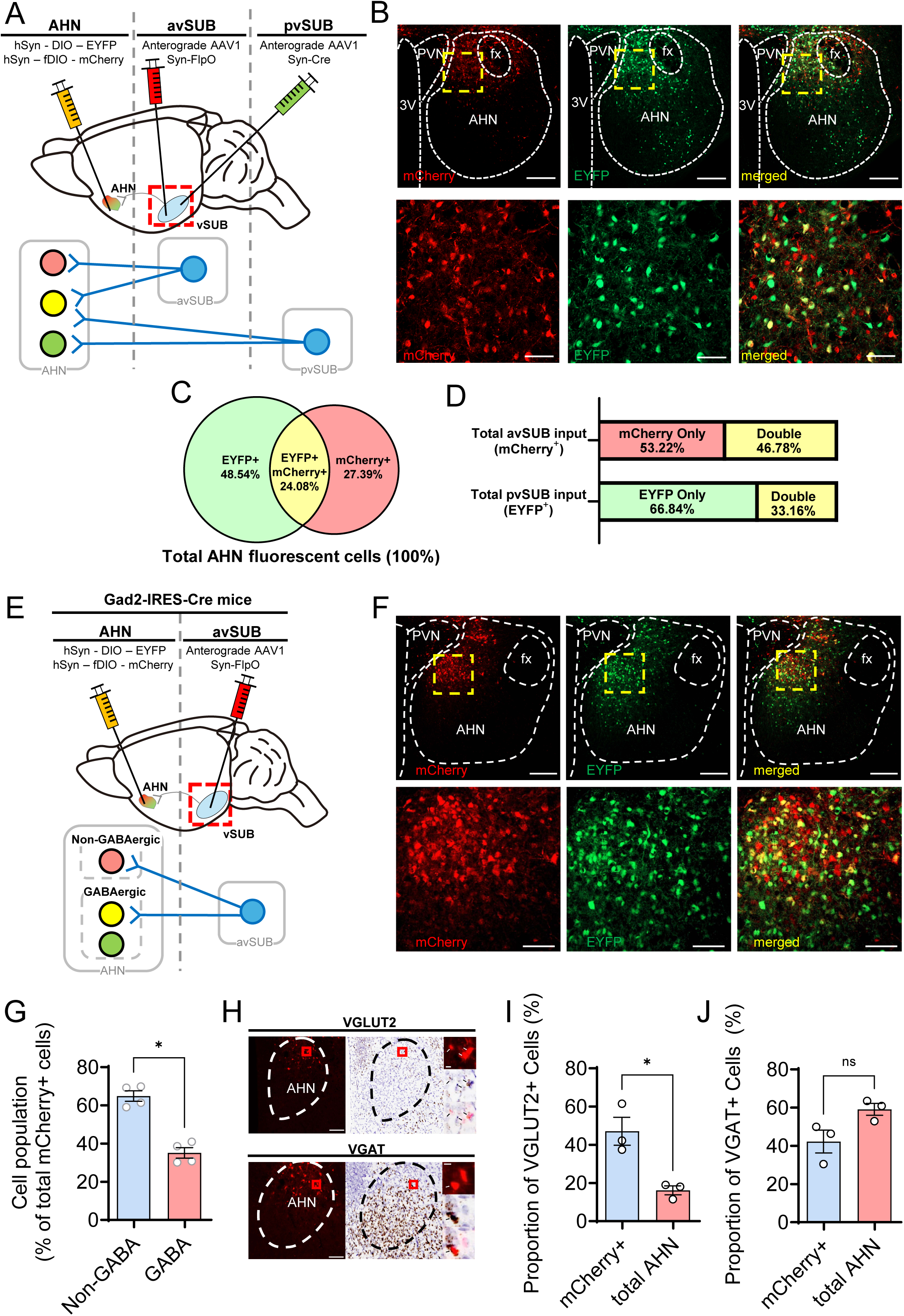
The anterior and posterior ventral subiculum (vSUB) target largely separate populations of anterior hypothalamic nucleus (AHN) neurons. (A) Timeline and strategy for dual-color anterograde transsynaptic tracing. AAV1-EF1a-FlpO was injected into the anterior ventral subiculum (avSUB) and AAV1-hSyn-Cre into the posterior ventral subiculum (pvSUB), together with Flp-dependent mCherry and Cre-dependent EYFP in the AHN. (B) Representative images of mCherry-labeled, EYFP-labeled and double-labeled AHN neurons. The paraventricular nucleus of the hypothalamus (PVN), the fornix (fx) and the third ventricle (3V) are shown as anatomical landmarks, and arrowheads mark double-labeled cells. (C) Proportions of AHN neurons receiving avSUB input alone, pvSUB input alone, or both, expressed as a percentage of all labeled AHN neurons (n = 4 mice). (D) The same data expressed separately for each input, as the percentage of avSUB-recipient or pvSUB-recipient neurons that were single or double labeled (n = 4 mice). (E) Strategy for identifying the transmitter phenotype of avSUB-recipient AHN neurons in Gad2-IRES-Cre mice, in which mCherry reports avSUB input and EYFP reports GABAergic identity. (F) Representative images, in which the PVN and the fx are shown as anatomical landmarks. Arrowheads mark double-labeled cells. (G) Proportions of non-GABAergic and GABAergic cells among avSUB-recipient AHN neurons (non-GABAergic versus GABAergic, p =.012, n = 4 mice). (H) Representative images of RNAscope in situ hybridization. mCherry fluorescence (left) and 3,3′-diaminobenzidine (DAB) labeling with hematoxylin counterstaining (right) were aligned, and the insets show a magnified double-labeled cell. (I) Proportion of VGLUT2-positive cells among avSUB-recipient AHN neurons and among all AHN neurons within the sampled regions of interest (p =.034, n = 3 mice). (J) Proportion of VGAT-positive cells in the same comparison (p =.167, n = 3 mice). Data are mean ± SEM. Panels G, I and J were analyzed by paired t test within each mouse. Statistics are reported in Table S1. Scale bars, 200 μm in the upper images and 50 μm in the magnified images of (B) and (F), 100 μm in (H), and 10 μm in the insets.

Given that the pvSUB preferentially targets AHN GABAergic neurons (15), we next characterized the postsynaptic cell types of the avSUB pathway. To achieve this, we injected AAV1-EF1a-FlpO into the avSUB and Flp-dependent mCherry and Cre-dependent EYFP into the AHN in Gad2-IRES-Cre mice (48), so that mCherry reported avSUB input and EYFP reported GABAergic identity (Figure 6E, 6F). Approximately two thirds of avSUB-recipient AHN neurons were non-GABAergic (Figure 6G). In situ hybridization supported this conclusion. Among avSUB-recipient AHN neurons, 47.1% were VGLUT2-positive, which was higher than the 16.2% of VGLUT2-positive neurons in the AHN as a whole and corresponds to an approximately threefold enrichment (Figure 6H, 6I). By contrast, the proportion of VGAT-positive cells among avSUB-recipient neurons did not differ from that in the AHN as a whole (Figure 6J). Taken together, these results show that the avSUB targets a predominantly non-GABAergic population of AHN neurons, which is anatomically distinct from the pvSUB-targeted population.

### avSUB→AHN PATHWAY MEDIATES SOCIAL BEHAVIOR WHEREAS pvSUB→AHN PATHWAY DRIVES ANXIETY-LIKE BEHAVIOR

We next asked whether the two pathways also differ functionally. Given that the pvSUB→AHN has been previously implicated in generalized anxiety (15), we first evaluated the behavioral role of both pathways using the OFT and the EPM. We expressed mCherry, hM3D(Gq), or hM4D(Gi) specifically in the postsynaptic AHN neurons targeted by either subregion (Figure 7A).

**Figure 7.**
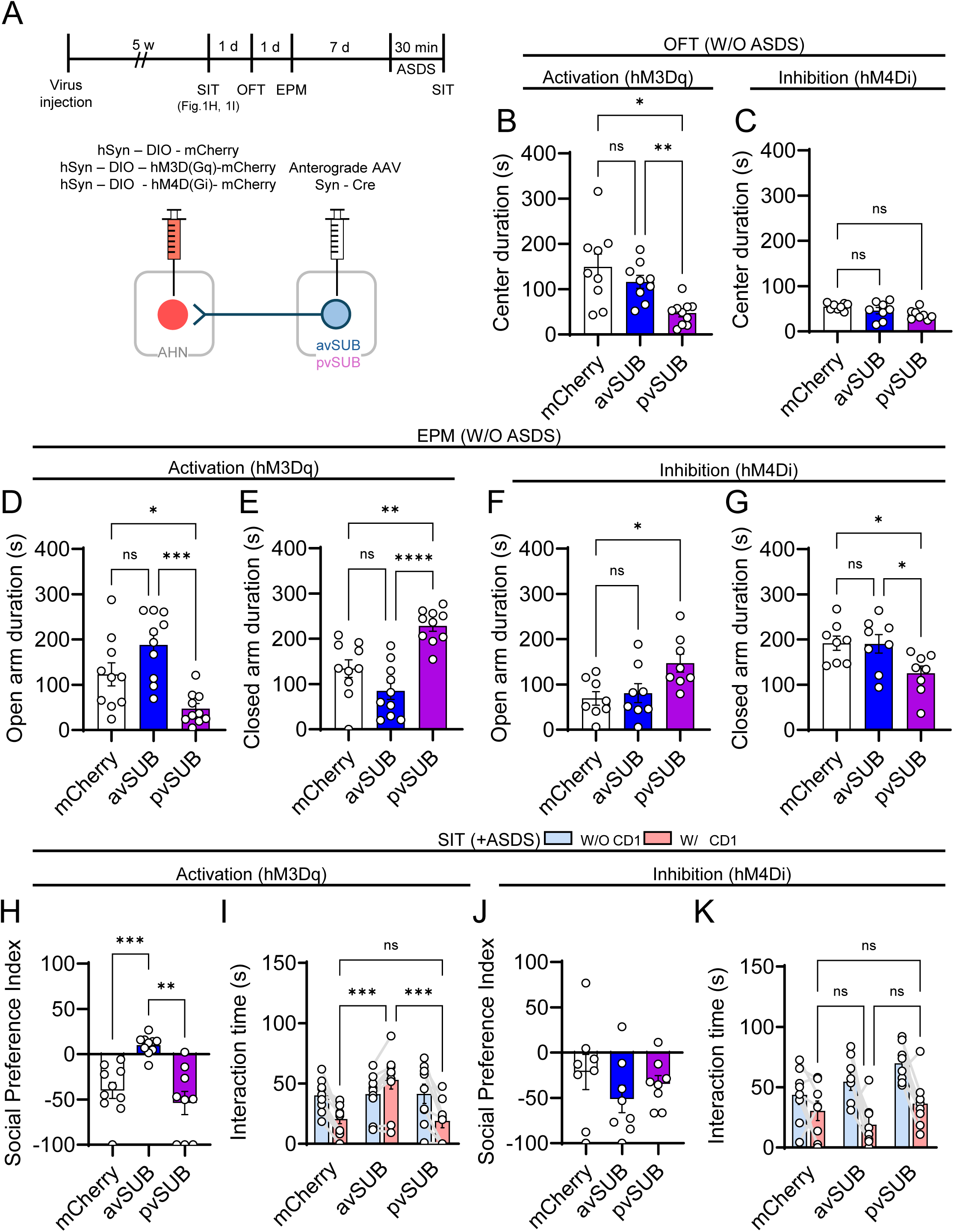
The anterior ventral subiculum (avSUB)→anterior hypothalamic nucleus (AHN) pathway controls social approach and the posterior ventral subiculum (pvSUB)→AHN pathway controls anxiety-like behavior. (A) Strategy and timeline for chemogenetic manipulation of AHN neurons receiving avSUB or pvSUB input. Anterograde transsynaptic AAV1-hSyn-Cre was injected into the avSUB or the pvSUB and Cre-dependent hM3D(Gq), hM4D(Gi), or mCherry into the AHN. Clozapine N-oxide (CNO) was given 30 min before testing in the stress-naive open field test (OFT) and elevated plus maze (EPM) experiments and 30 min before the stressor in the social interaction test (SIT). (B) Time spent in the center of the open field in stress-naive mice after activation with hM3D(Gq) (mCherry versus avSUB p =.667, mCherry versus pvSUB p =.021, avSUB versus pvSUB p =.003, n = 9, 9 and 10 mice). (C) Time spent in the center of the open field in stress-naive mice after inhibition with hM4D(Gi) (no effect of group, n = 8 mice per group). (D) Open arm duration in the EPM in stress-naive mice after activation with hM3D(Gq) (mCherry versus avSUB p =.088, mCherry versus pvSUB p =.041, avSUB versus pvSUB p <.001, n = 10 mice per group). (E) Closed arm duration in the same mice (mCherry versus avSUB p =.121, mCherry versus pvSUB p =.002, avSUB versus pvSUB p <.0001, n = 10 mice per group). (F) Open arm duration in the EPM in stress-naive mice after inhibition with hM4D(Gi) (mCherry versus avSUB p =.900, mCherry versus pvSUB p =.021, n = 8 mice per group). (G) Closed arm duration in the same mice (mCherry versus avSUB p =.998, mCherry versus pvSUB p =.037, avSUB versus pvSUB p =.042, n = 8 mice per group). (H) Social preference index after acute social defeat stress (ASDS) with hM3D(Gq) (mCherry versus avSUB p <.001, avSUB versus pvSUB p =.003, n = 10, 10 and 9 mice). (I) Interaction time after ASDS with hM3D(Gq) (with the target present, mCherry versus avSUB p <.001, mCherry versus pvSUB not significant, avSUB versus pvSUB p <.001, n = 10, 10 and 9 mice). (J) Social preference index after ASDS with hM4D(Gi) (all comparisons not significant, n = 8 mice per group). (K) Interaction time after ASDS with hM4D(Gi) (all comparisons not significant, n = 8 mice per group). In panels I and K, light blue bars show the epoch without a target and light red bars the epoch with a target. Data are mean ± SEM. Panels B, H and J were analyzed by Brown-Forsythe and Welch ANOVA with Dunnett’s T3 multiple comparisons test, panel C by one-way ANOVA, panels D to G by one-way ANOVA with Tukey’s multiple comparisons test, and panels I and K by two-way repeated measures ANOVA with Tukey’s multiple comparisons test. Statistics are reported in Table S1.

Chemogenetic activation of the pvSUB→AHN pathway significantly reduced center duration in the OFT (Figure 7B) and time spent in the open arms of the EPM (Figure 7D), and increased closed arm duration (Figure 7E), indicating a significant increase in anxiety-like behavior. Conversely, inhibition of the pvSUB→AHN pathway increased open arm time and reduced closed arm duration in the EPM (Figure 7F, 7G), consistent with an anxiolytic effect, although OFT performance remained largely unaltered (Figure 7C). In contrast, bidirectional manipulation of the avSUB→AHN pathway had no effect on anxiety-like behavior compared to mCherry controls (Figure 7B to 7G). These findings demonstrate that the pvSUB→AHN pathway is tightly coupled to anxiety-like behavior, whereas the avSUB→AHN pathway is not.

We next examined their respective roles in ASDS-induced social avoidance. As we already found in Figure 4G, chemogenetic activation of the avSUB→AHN pathway attenuated ASDS-induced social avoidance. However, activation of the pvSUB→AHN pathway did not, with mice performing indistinguishably from mCherry controls (Figure 7H, 7I). Conversely, inhibition of either pathway failed to alter social interaction following ASDS (Figure 7J, 7K). For the avSUB→AHN pathway, this lack of effect is attributable to a floor effect as ASDS already suppresses social interaction. For the pvSUB→AHN pathway, this result aligns with the absence of any behavioral effect following identical manipulation in stress-naive mice (Figure 1I). Collectively, we conclude that the avSUB→AHN and pvSUB→AHN pathways serve distinct behavioral roles, driving social approach and generalized anxiety, respectively.

## DISCUSSION

Studies on social defeat stress have primarily emphasized neural adaptations that accumulate across repeated defeat or persist after the defeat has ceased. However, much less is known about the circuit and synaptic events occurring during the social defeat itself that contribute to the emergence of social avoidance. In this study, we demonstrate that ASDS induced an immediate and pathway-specific synaptic depression in the avSUB→AHN circuit, establishing a direct link between this plasticity and social avoidance. A single defeat episode selectively decreased presynaptic release probability at avSUB→AHN synapses, while sparing the pvSUB→AHN pathway. This depression was strictly dependent on glucocorticoid receptor and CB1R signaling. Crucially, local CB1R blockade within the AHN attenuated both the synaptic depression and the behavioral avoidance. Moreover, silencing the AHN neurons that receive avSUB input was sufficient to induce social avoidance in the absence of any stressor. Finally, we establish a striking anatomical and functional dichotomy in which the avSUB→AHN pathway mediates social approach whereas the pvSUB→AHN pathway exclusively governs anxiety-like behavior. This highlights a highly specialized, parallel processing architecture within the vSUB for coping with distinct emotional threats.

A critical insight from our data is that the synaptic depression in the avSUB→AHN pathway was transient, fully recovering within 24 hours, whereas the behavioral avoidance was long-lasting. To resolve this temporal contradiction, we can compare the outcomes of our chemogenetic and optogenetic manipulations. Chemogenetic activation of the avSUB→AHN pathway during ASDS conferred long-lasting protection, preventing social avoidance at both 30-minute and 24-hour time points. In contrast, optogenetic activation applied after ASDS provided only a transient, light-locked rescue, conferring no enduring protection and leaving the avoidance fully intact at 24 hours. These findings suggest that acute suppression of the avSUB→AHN circuit acts as an initial trigger rather than a chronic maintainer of social avoidance. Maintaining circuit tone during ASDS blocks the transition into a persistent avoidance state, whereas activating the pathway after that state is established overrides the avoidance only while the light is on. Thus, the transient synaptic depression of the avSUB→AHN pathway might serve as a critical, time-sensitive gateway that propagates into long-lasting behavioral adaptations.

While our findings establish the avSUB→AHN pathway as a critical mediator of acute social avoidance, we acknowledge that the prevailing literature on social defeat has heavily emphasized the nucleus accumbens (NAc) and the mesolimbic dopamine system in the emergence of depressive-like behaviors and social withdrawal (49). This divergence likely reflects the distinction between acute trauma and chronic load, because the classical NAc-dependent models describe maladaptive plasticity accumulating over days of repeated defeat whereas our study isolates the immediate circuitry that translates a single episode into an acute behavioral change. Therefore, it is plausible that the rapid eCB-dependent gating in the avSUB→AHN circuit initiates the primary adaptive response to acute social threat, which may subsequently recruit downstream structures, including the NAc, to consolidate a long-term depressive phenotype during chronic stress.

To date, vSUB has been broadly implicated in emotional regulation and stress responses. Previous studies have established that a vSUB input to AHN GABAergic neurons drives innate, nonsocial avoidance (15). Crucially, they reported that this specific projection is not engaged while mice investigate a conspecific, raising the question of how the vSUB contributes to social paradigms. Our current findings unveiled a highly specialized, topographically organized functional dichotomy within the vSUB. We demonstrate that the vSUB has largely segregated parallel circuits. The previous findings (15) likely reflect the activity of the posterior population (pvSUB→AHN), which our data confirm is completely uncoupled from social interaction but exclusively governs generalized, non-social anxiety-like behaviors. In contrast, we identified an anatomically distinct anterior projection (avSUB→AHN) that serves a fundamentally different purpose. These circuits are also differentially engaged by acute social defeat, and their artificial activation drives opposing behavioral states. While stimulating the pvSUB→AHN pathway exacerbates non-social anxiety, activation of the avSUB→AHN pathway strongly promotes social approach and rescues ASDS-induced social avoidance. Furthermore, our dual-transsynaptic tracing and in situ hybridization data reveal that while the avSUB and pvSUB pathways exhibit some convergence, they preferentially innervate largely distinct neuronal subpopulations within the AHN with differing GABAergic profiles. This topographical and functional double dissociation demonstrates that the vSUB does not act as a single, uniform stress center. Instead, it uses separate parallel circuits to process different types of stressor independently.

The two target populations are largely, though not entirely, distinct. About a quarter of labeled AHN neurons received both avSUB and pvSUB inputs. Moreover, because the two injection sites lie close together along the anteroposterior axis, viral spread into the intervening subiculum likely inflates this overlap fraction. Because overlap would dilute rather than produce a functional divergence, the dissociation we observed is likely a conservative underestimate.

The anatomical and functional divergence we describe must ultimately rest on a difference between the avSUB and pvSUB projection neurons themselves. However, what inherently differentiates them remains an open question. While our current data do not provide an answer, we speculate that this functional dichotomy arises because the vSUB and ventral hippocampus do not form a single uniform structure. Rather, their molecular properties gradually change along the anterior-posterior axis (18,50). Interestingly, vSUB neurons expressing dopamine D1 and D2 receptors form distinct populations with opposing behavioral roles, driving anxiety-like avoidance and approach, respectively (51). Because this aligns closely with our functional findings, we hypothesize that the avSUB and pvSUB projection neurons might correspond to these D2 and D1 subpopulations, respectively. As previous studies have not linked receptor identity to specific efferent targets or anteroposterior location, mapping D1 and D2 receptor expression onto our identified pathways would be a critical next step to resolve the molecular underpinnings of this circuit.

Finally, our study elucidates the precise synaptic mechanism by which ASDS is translated into lasting social avoidance. Glucocorticoid-driven, CB1R-mediated suppression of synaptic transmission is a well-established mechanism at hippocampal, hypothalamic and amygdalar synapses, where it finely tunes neuroendocrine and emotional outputs (22,30–32,52–55). Our findings extend this mechanism in two ways. We place it on a long-range hippocampal input to the hypothalamus rather than on a local hypothalamic synapse, and we attach it to a social behavioral readout rather than to a neuroendocrine one. We demonstrate that ASDS induces a rapid, CB1R-dependent presynaptic depression specifically at avSUB→AHN terminals. Importantly, local infusion of the CB1R antagonist AM251 during stress successfully blocked both the synaptic depression and the resulting social avoidance. These findings point to the eCB-CB1R signaling as a critical mediator for stress-induced social deficits, highlighting a novel therapeutic target for reversing maladaptive social avoidance.

## Supporting information

Supplemetary methods and materials, supplementary figures, statistic table

## ACKNOWLEDGMENTS AND DISCLOSURES

C.W. led the project and performed stereotaxic surgery, viral tracing, social defeat and behavioral testing, pathway-specific chemogenetic and optogenetic manipulation, slice electrophysiology and pharmacology, and histology and imaging, and analyzed the data. S.L., a co-first author, performed the behavioral and slice electrophysiology experiments together with C.W., including intra-AHN drug infusion and systemic pharmacology, and contributed equally to the design and analysis of those experiments. C.L. contributed to optic fiber implantation and in vivo optogenetic stimulation and to slice electrophysiology and pharmacology. D.Y. performed the RNAscope in situ hybridization and the atlas registration, supervised by S.C., who also provided the resources for that work. S.J.K., K.C. and S.C. contributed to data analysis and study interpretation. K.S.S. conceived and supervised the project, and C.W. and K.S.S. acquired funding. C.W., S.L., K.C. and K.S.S. wrote the original draft, and all authors contributed to article editing and figure design.

This work was supported by the National Research Foundation of Korea (Grant No. 2020R1A2C1011434 and RS-2025-16070398 to K.S.S., and RS-2025-25424624 to C.W.). The authors report no biomedical financial interests or potential conflicts of interest.

During the preparation of this work, the authors used Claude (Opus5.0, Anthropic) to improve grammar and language clarity. After using this tool, the authors reviewed and edited the content as needed and take full responsibility for the content of the published article.

## REFERENCES

1. Gellner AK, Voelter J, Schmidt U, Beins EC, Stein V, Philipsen A, Hurlemann R (2021): Molecular and neurocircuitry mechanisms of social avoidance. Cell Mol Life Sci 78: 1163–1189.

2. Golden SA, Covington HE 3rd, Berton O, Russo SJ (2011): A standardized protocol for repeated social defeat stress in mice. Nat Protoc 6: 1183–1191.

3. Krishnan V, Han MH, Graham DL, Berton O, Renthal W, Russo SJ, et al. (2007): Molecular adaptations underlying susceptibility and resistance to social defeat in brain reward regions. Cell 131: 391–404.

4. Berton O, McClung CA, DiLeone RJ, Krishnan V, Renthal W, Russo SJ, et al. (2006): Essential role of BDNF in the mesolimbic dopamine pathway in social defeat stress. Science 311: 864–868.

5. Radzicki D, McCann KE, Alexander GM, Dudek SM (2025): Hippocampal area CA2 activity supports social investigation following an acute social stress. Mol Psychiatry 30: 2284–2296.

6. Herman JP, Mueller NK (2006): Role of the ventral subiculum in stress integration. Behav Brain Res 174: 215–224.

7. Herman JP, Dolgas CM, Carlson SL (1998): Ventral subiculum regulates hypothalamo-pituitary-adrenocortical and behavioural responses to cognitive stressors. Neuroscience 86: 449–459.

8. Miller CN, Li Y, Beier KT, Aoto J (2025): Acute stress causes sex-specific changes to ventral subiculum synapses, circuitry, and anxiety-like behavior. Nat Commun 16: 5604.

9. Kemp GM, Altimimi HF, Nho Y, Heir R, Klyczek A, Stellwagen D (2022): Sustained TNF signaling is required for the synaptic and anxiety-like behavioral response to acute stress. Mol Psychiatry 27: 4474–4484.

10. Canteras NS (2002): The medial hypothalamic defensive system: Hodological organization and functional implications. Pharmacol Biochem Behav 71: 481–491.

11. Gross CT, Canteras NS (2012): The many paths to fear. Nat Rev Neurosci 13: 651–658.

12. Xie Z, Gu H, Huang M, Cheng X, Shang C, Tao T, et al. (2022): Mechanically evoked defensive attack is controlled by GABAergic neurons in the anterior hypothalamic nucleus. Nat Neurosci 25: 72–85.

13. Bang JY, Zhao J, Rahman M, St-Cyr S, McGowan PO, Kim JC (2022): Hippocampus–anterior hypothalamic circuit modulates stress-induced endocrine and behavioral response. Front Neural Circuits 16: 894722.

14. Hong CY, Din JS, Chang H, Bang JY, Kim JC (2025): Anterior hypothalamic nucleus drives distinct defensive responses through cell-type-specific activity. iScience 28: 112097.

15. Yan JJ, Ding XJ, He T, Chen AX, Zhang W, Yu ZX, et al. (2022): A circuit from the ventral subiculum to anterior hypothalamic nucleus GABAergic neurons essential for anxiety-like behavioral avoidance. Nat Commun 13: 7464.

16. Canteras NS, Swanson LW (1992): Projections of the ventral subiculum to the amygdala, septum, and hypothalamus: A PHAL anterograde tract-tracing study in the rat. J Comp Neurol 324: 180–194.

17. Cembrowski MS, Phillips MG, DiLisio SF, Shields BC, Winnubst J, Chandrashekar J, et al. (2018): Dissociable structural and functional hippocampal outputs via distinct subiculum cell classes. Cell 173: 1280–1292.e18.

18. Ding SL, Yao Z, Hirokawa KE, Nguyen TN, Graybuck LT, Fong O, et al. (2020): Distinct transcriptomic cell types and neural circuits of the subiculum and prosubiculum along the dorsal–ventral axis. Cell Rep 31: 107648.

19. Toth I, Neumann ID, Slattery DA (2012): Social fear conditioning: A novel and specific animal model to study social anxiety disorder. Neuropsychopharmacology 37: 1433–1443.

20. Xu H, Liu L, Tian Y, Wang J, Li J, Zheng J, et al. (2019): A disinhibitory microcircuit mediates conditioned social fear in the prefrontal cortex. Neuron 102: 668–682.e5.

21. Marsh JS, Teixeira C, Gavade S, Johnston C, Baranwal S, Snyder CN, et al. (2025): Ventral subiculum control of avoidance behavior and hypothalamic-pituitary-adrenal axis reactivity via the bed nucleus of the stria terminalis in male and female mice. Psychoneuroendocrinology 171: 107229.

22. Karst H, Berger S, Turiault M, Tronche F, Schütz G, Joëls M (2005): Mineralocorticoid receptors are indispensable for nongenomic modulation of hippocampal glutamate transmission by corticosterone. Proc Natl Acad Sci U S A 102: 19204–19207.

23. Wamsteeker JI, Bains JS (2010): A synaptocentric view of the neuroendocrine response to stress. Eur J Neurosci 32: 2011–2021.

24. Kuzmiski JB, Marty V, Baimoukhametova DV, Bains JS (2010): Stress-induced priming of glutamate synapses unmasks associative short-term plasticity. Nat Neurosci 13: 1257–1264.

25. Chevaleyre V, Takahashi KA, Castillo PE (2006): Endocannabinoid-mediated synaptic plasticity in the CNS. Annu Rev Neurosci 29: 37–76.

26. Castillo PE, Younts TJ, Chávez AE, Hashimotodani Y (2012): Endocannabinoid signaling and synaptic function. Neuron 76: 70–81.

27. Kano M, Ohno-Shosaku T, Hashimotodani Y, Uchigashima M, Watanabe M (2009): Endocannabinoid-mediated control of synaptic transmission. Physiol Rev 89: 309–380.

28. Gerdeman GL, Ronesi J, Lovinger DM (2002): Postsynaptic endocannabinoid release is critical to long-term depression in the striatum. Nat Neurosci 5: 446–451.

29. Robbe D, Kopf M, Remaury A, Bockaert J, Manzoni OJ (2002): Endogenous cannabinoids mediate long-term synaptic depression in the nucleus accumbens. Proc Natl Acad Sci U S A 99: 8384–8388.

30. Di S, Malcher-Lopes R, Halmos KC, Tasker JG (2003): Nongenomic glucocorticoid inhibition via endocannabinoid release in the hypothalamus: A fast feedback mechanism. J Neurosci 23: 4850–4857.

31. Di S, Malcher-Lopes R, Marcheselli VL, Bazan NG, Tasker JG (2005): Rapid glucocorticoid-mediated endocannabinoid release and opposing regulation of glutamate and γ-aminobutyric acid inputs to hypothalamic magnocellular neurons. Endocrinology 146: 4292–4301.

32. Di S, Maxson MM, Franco A, Tasker JG (2009): Glucocorticoids regulate glutamate and GABA synapse-specific retrograde transmission via divergent nongenomic signaling pathways. J Neurosci 29: 393–401.

33. Patel S, Roelke CT, Rademacher DJ, Hillard CJ (2005): Inhibition of restraint stress-induced neural and behavioural activation by endogenous cannabinoid signalling. Eur J Neurosci 21: 1057–1069.

34. Di S, Itoga CA, Fisher MO, Solomonow J, Roltsch EA, Gilpin NW, Tasker JG (2016): Acute stress suppresses synaptic inhibition and increases anxiety via endocannabinoid release in the basolateral amygdala. J Neurosci 36: 8461–8470.

35. Droste SK, de Groote L, Atkinson HC, Lightman SL, Reul JMHM, Linthorst ACE (2008): Corticosterone levels in the brain show a distinct ultradian rhythm but a delayed response to forced swim stress. Endocrinology 149: 3244–3253.

36. Loomba N, Patel S (2025): Circuit mechanisms governing endocannabinoid modulation of affective behaviour and stress adaptation. Nat Rev Neurosci 26: 677–697.

37. Wei D, Allsop S, Tye K, Piomelli D (2017): Endocannabinoid signaling in the control of social behavior. Trends Neurosci 40: 385–396.

38. Litvin Y, Phan A, Hill MN, Pfaff DW, McEwen BS (2013): CB1 receptor signaling regulates social anxiety and memory. Genes Brain Behav 12: 479–489.

39. Beins EC, Beiert T, Jenniches I, Hansen JN, Leidmaa E, Schrickel JW, Zimmer A (2021): Cannabinoid receptor 1 signalling modulates stress susceptibility and microglial responses to chronic social defeat stress. Transl Psychiatry 11: 164.

40. Zingg B, Chou XL, Zhang ZG, Mesik L, Liang F, Tao HW, Zhang LI (2017): AAV-mediated anterograde transsynaptic tagging: Mapping corticocollicular input-defined neural pathways for defense behaviors. Neuron 93: 33–47.

41. Zingg B, Peng B, Huang J, Tao HW, Zhang LI (2020): Synaptic specificity and application of anterograde transsynaptic AAV for probing neural circuitry. J Neurosci 40: 3250–3267.

42. Armbruster BN, Li X, Pausch MH, Herlitze S, Roth BL (2007): Evolving the lock to fit the key to create a family of G protein-coupled receptors potently activated by an inert ligand. Proc Natl Acad Sci U S A 104: 5163–5168.

43. Klapoetke NC, Murata Y, Kim SS, Pulver SR, Birdsey-Benson A, Cho YK, et al. (2014): Independent optical excitation of distinct neural populations. Nat Methods 11: 338–346.

44. Zucker RS, Regehr WG (2002): Short-term synaptic plasticity. Annu Rev Physiol 64: 355–405.

45. Tervo DGR, Hwang BY, Viswanathan S, Gaj T, Lavzin M, Ritola KD, et al. (2016): A designer AAV variant permits efficient retrograde access to projection neurons. Neuron 92: 372–382.

46. Gomez JL, Bonaventura J, Lesniak W, Mathews WB, Sysa-Shah P, Rodriguez LA, et al. (2017): Chemogenetics revealed: DREADD occupancy and activation via converted clozapine. Science 357: 503–507.

47. Kitanishi T, Tashiro M, Kitanishi N, Mizuseki K (2022): Intersectional, anterograde transsynaptic targeting of neurons receiving monosynaptic inputs from two upstream regions. Commun Biol 5: 149.

48. Taniguchi H, He M, Wu P, Kim S, Paik R, Sugino K, et al. (2011): A resource of Cre driver lines for genetic targeting of GABAergic neurons in cerebral cortex. Neuron 71: 995–1013.

49. Russo SJ, Nestler EJ (2013): The brain reward circuitry in mood disorders. Nat Rev Neurosci 14: 609–625.

50. Cembrowski MS, Bachman JL, Wang L, Sugino K, Shields BC, Spruston N (2016): Spatial gene-expression gradients underlie prominent heterogeneity of CA1 pyramidal neurons. Neuron 89: 351–368.

51. Godino A, Salery M, Minier-Toribio AM, Patel V, Fullard JF, Kondev V, et al. (2025): Dopamine D1–D2 signalling in hippocampus arbitrates approach and avoidance. Nature 643: 448–457.

52. Patel S, Kingsley PJ, Mackie K, Marnett LJ, Winder DG (2009): Repeated homotypic stress elevates 2-arachidonoylglycerol levels and enhances short-term endocannabinoid signaling at inhibitory synapses in basolateral amygdala. Neuropsychopharmacology 34: 2699–2709.

53. Herman JP, Tasker JG (2016): Paraventricular hypothalamic mechanisms of chronic stress adaptation. Front Endocrinol (Lausanne) 7: 137.

54. Karst H, Berger S, Erdmann G, Schütz G, Joëls M (2010): Metaplasticity of amygdalar responses to the stress hormone corticosterone. Proc Natl Acad Sci U S A 107: 14449–14454.

55. Ramikie TS, Nyilas R, Bluett RJ, Gamble-George JC, Hartley ND, Mackie K, et al. (2014): Multiple mechanistically distinct modes of endocannabinoid mobilization at central amygdala glutamatergic synapses. Neuron 81: 1111–1125.

