## Supplementary material for "Acute social defeat induces cannabinoid type 1 receptor-dependent synaptic depression in the anterior ventral subiculum to anterior hypothalamic nucleus pathway to drive social avoidance in male mice": Supplemetary methods and materials, supplementary figures, statistic table

### CONTENTS

| | |
|---|---|
| Supplemental Methods and Materials | 2 |
| Supplementary Figures S1 to S5 with legends | 11 |
| Supplemental Table S1. Statistical analyses | 21 |

### SUPPLEMENTAL METHODS AND MATERIALS

#### Animals

Adult male wild-type C57BL/6N mice (6–12 weeks of age; Orient Bio) and male Gad2-IRES-Cre mice (B6N.Cg-Gad2tm2(cre)Zjh/J; Strain #019022, The Jackson Laboratory; 12–16 weeks of age) were used, and male CD-1 retired breeder mice (6 months of age; DBL) served as aggressors. Mice were group-housed on a 12-h light/dark cycle at 20–24°C and 30–60% humidity with food and water available ad libitum, and were single-housed for 1 day after stereotaxic surgery.

#### Stereotaxic Surgery and Viral Vectors

Mice were anesthetized with isoflurane and given bupivacaine (5 mg/kg) for analgesia. Surgery was performed on a stereotaxic frame (Stoelting), and viruses were delivered with a Picospritzer III (Parker) at a slow rate (50 nL/min). Coordinates relative to bregma were: anterior ventral subiculum (avSUB), anteroposterior (AP) −3.50 mm, mediolateral (ML) ±2.90 mm, dorsoventral (DV) −4.95 mm; posterior ventral subiculum (pvSUB), AP −4.15 mm, ML ±3.00 mm, DV −4.60 mm; anterior hypothalamic nucleus (AHN), AP −0.82 mm, ML ±0.50 mm, DV −4.95 mm. Mice recovered in their home cage for at least 5 weeks before experiments.

For optogenetic experiments, optic fibers (1.25 mm ceramic ferrule, 200 μm core, numerical aperture 0.37, 4.9 mm length; Inper) were implanted bilaterally over the AHN at a 20° angle to avoid damage to the midline, 1 week before behavioral testing, and secured with dental cement. For local drug infusion, a guide cannula (C235GS-5-0.6/SPC; P1 Technologies) was implanted bilaterally over the AHN 3 weeks after virus injection and 2 weeks before testing.

The following viral vectors were used: AAV1-hSyn-Cre (pENN.AAV.hSyn.Cre.WPRE.hGH; Addgene; titer ≥1 × $10^{13}$ vg/mL), AAV1-EF1a-Flpo (Addgene; ≥7 × $10^{12}$ vg/mL), AAVretro-EF1a-Cre (Addgene; ≥7 × $10^{12}$ vg/mL), AAVretro-CAG-Cre (UNC Vector Core; ≥7 × $10^{12}$ vg/mL), AAV2-hSyn-DIO-hM3D(Gq)-mCherry (Addgene; ≥6 × $10^{12}$ vg/mL), AAV2-hSyn-DIO-hM4D(Gi)-mCherry (Addgene; ≥5 × $10^{12}$ vg/mL), AAV9-hSyn-DIO-mCherry (Addgene; ≥1 × $10^{13}$ vg/mL), AAV9-Ef1a-fDIO-mCherry (Addgene; ≥1 × $10^{13}$ vg/mL), AAV5-Ef1a-DIO-EYFP (Addgene; ≥1 × $10^{13}$ vg/mL), AAV9-Syn-ChrimsonR-tdTomato (Addgene; ≥1 × $10^{13}$ vg/mL), AAV8-hSyn-FLEX-Chronos-GFP (UNC Vector Core; ≥7 × $10^{12}$ vg/mL), and AAV5-hSyn-Chronos-GFP (UNC Vector Core; ≥7 × $10^{12}$ vg/mL). Vectors obtained from the UNC Vector Core were produced from materials provided by E.S. Boyden (Massachusetts Institute of Technology). Injection volumes were 0.5 μL per site in the avSUB, 0.5–0.6 μL in the pvSUB, and 0.4–0.5 μL in the AHN.

To visualize the two subicular populations in the same brain, AAV-hSyn-Chronos-GFP was injected into the avSUB and AAV-Syn-ChrimsonR-tdTomato into the pvSUB.

For slice recordings, the Cre-independent AAV5-hSyn-Chronos-GFP was injected into the avSUB or the pvSUB and light-evoked currents were recorded from AHN neurons. For in vivo optogenetics, AAVretro-CAG-Cre was injected into the AHN together with Cre-dependent AAV8-hSyn-FLEX-Chronos-GFP, or a Cre-dependent GFP control, into the avSUB, so that Chronos was expressed selectively in AHN-projecting avSUB neurons and their terminals could be stimulated through the fiber implanted over the AHN. For the chemogenetic manipulation of AHN-projecting avSUB neurons, AAVretro-EF1a-Cre was injected into the AHN together with Cre-dependent DREADDs or mCherry into the avSUB. For chemogenetic control of the AHN neurons receiving subicular input, anterograde transsynaptic AAV1-hSyn-Cre was injected into the avSUB or the pvSUB together with Cre-dependent DREADDs or mCherry into the AHN.

### Stress Protocols

CD-1 retired breeders were screened as aggressors when they attacked a C57BL/6N screener mouse (7–8 weeks of age) within 1 min on three consecutive screening days. Test mice were 11–12 weeks of age at the time of defeat. A different aggressor was used for each session.

Subthreshold social defeat stress (SSDS) consisted of 5 min of physical defeat followed by 25 min in the animal’s own home cage, and therefore was both shorter than acute social defeat stress (ASDS) and lacked the sensory exposure phase.

### Behavioral Tests

All behavioral experiments were conducted in a soundproof chamber. Mice were transferred from the housing facility and acclimated for 1 h before testing. Movement was recorded and analyzed with EthoVision 3.0 (Noldus), and all apparatus was cleaned with 70% ethanol between subjects and trials. All behavioral measures were acquired and scored automatically by the video-tracking software, so no manual scoring was involved.

### Social Interaction Test

The social interaction test (SIT) used an open-field box (40 × 40 × 30 cm) with a perforated Plexiglas target cage (8 × 6 × 30 cm) centered against one wall, and the interaction zone extended 8 cm around the cage. Mice were placed in the center and allowed to explore for 2 min 30 s in the absence and then in the presence of an unfamiliar CD-1 target. Where a second test was performed 24 hours later, a different CD-1 target was used. The social interaction ratio was calculated as the interaction time with the target divided by the interaction time without the target, and the social preference index (SPI) as

$$SPI = [(t_1 - t_0) / (t_1 + t_0)] \times 100,$$

where $t_1$ and $t_0$ are the interaction times in the presence and absence of the target, respectively.

**Light-Stimulation Social Interaction Test**

After the standard phases without and with the CD-1 target (2 min 30 s each), the session continued for three further cycles in which light was turned on and off every 2 min 30 s with the target still present. Blue light (470 nm, 15 mW at the fiber tip, 5-ms pulses at 20 Hz) was delivered throughout each 2 min 30 s light-on epoch from a diode laser (MDL-III-470 with a PSU-III-LED power supply; CNI Laser). Mice were tested first in the unstressed state and again 1 week later at 30 min and 24 hours after ASDS, with a different aggressor and a different target for each session.

**Open Field Test**

Mice were left undisturbed in the experimental room for about 10 min, then placed in the center (20 × 20 cm) of a novel open-field arena (40 × 40 × 30 cm) under dim light (40 lx) and allowed to explore for 10 min. Time spent in and entries into the center were recorded. The open field test was performed first and the elevated plus maze on the following day.

**Elevated Plus Maze**

The maze consisted of two open arms (30 × 5 × 2 cm) and two closed arms (30 × 5 × 30 cm) extending from a central area (10 × 10 cm), elevated 50 cm above the floor, under dim light (40 lx). Mice were placed in the center facing an open arm and allowed to explore freely for 5 min. Time spent in the open and closed arms and the number of arm entries were recorded.

### Drugs

CNQX (10 μM; Tocris) and D-APV (30 μM; Tocris) were bath-applied to confirm that light-evoked currents were glutamatergic. To test glucocorticoid involvement, slices from naive mice were pre-incubated in artificial cerebrospinal fluid (aCSF) containing corticosterone (100 nM; Sigma-Aldrich) for 30 min before recording, and RU486 (mifepristone, 20 mg/kg; Tocris) was injected intraperitoneally 30 min before ASDS. ACEA (5 μM; Tocris) and AM251 (5 μM for bath application; Tocris) were used to probe cannabinoid type 1 receptor (CB1R) signaling in slices. For local blockade, AM251 (0.3 mg/kg in 500 nL per hemisphere) was infused bilaterally into the AHN through the implanted cannula 30 min before ASDS. Because the ASDS protocol lasts 30 min, the SIT began 1 h after infusion. Clozapine-N-oxide (Tocris) was injected intraperitoneally in 100 μL 30 min before stress, at 1.5 mg/kg for hM3D(Gq) experiments and 5.0 mg/kg for hM4D(Gi) experiments, and 30 min before testing in experiments performed without a stressor. Vehicle was 5% dimethyl sulfoxide in phosphate-buffered saline, and control animals received an equal volume.

### Slice Preparation and Whole-Cell Recording

Mice were deeply anesthetized with isoflurane and transcardially perfused with 4°C aCSF containing (in mM): 124 NaCl, 2.5 KCl, 1.2 $NaH_2PO_4$, 24 $NaHCO_3$, 5 HEPES,

13 glucose, 2 $MgSO_4$, and 2 $CaCl_2$. Brains were rapidly removed and cut into 250-μm coronal sections on a vibratome (VT1000S, Leica) while submerged in 4°C aCSF. Slices were transferred to a 32°C recovery solution containing (in mM): 92 NMDG, 2.5 KCl, 1.2 $NaH_2PO_4$, 30 $NaHCO_3$, 20 HEPES, 25 glucose, 5 sodium ascorbate, 2 thiourea, 3 sodium pyruvate, 10 $MgSO_4$, and 0.5 $CaCl_2$ for 14 min, and were then held in aCSF at room temperature until recording.

Recordings were made in whole-cell voltage clamp at a holding potential of −70 mV, and the paired-pulse ratio was calculated as the amplitude of the second light-evoked current divided by that of the first, using a 100-ms interstimulus interval. Recordings were made with an EPC10 amplifier (HEKA) and sampled at 10 kHz. Pipettes pulled from borosilicate glass (Warner Instruments) had tip resistances of 4–5 MΩ when filled with internal solution containing (in mM): 130 Cs-methanesulfonate, 10 HEPES, 0.5 EGTA, 8 NaCl, 1 TEA-Cl, 10 $Na_2$-phosphocreatine, 1 QX-314, 4 Mg-ATP, 0.4 Na-GTP, and 0.1 spermine, adjusted to pH 7.2–7.3 with CsOH. Slices were superfused with 30°C aCSF at 1.6 mL/min and neurons were visualized with a 40× water-immersion objective (BX51WI, Olympus). Recordings were discarded if series resistance exceeded 20 MΩ or changed by more than 20%. Chronos-expressing terminals were stimulated with single 5-ms pulses of 470-nm light (10 mW) delivered through the objective from a 470-nm LED controlled by a T-Cube LED driver (LEDD1B, 1200 mA maximum drive current; Thorlabs). Data were analyzed with Patchmaster (HEKA) and Igor Pro 6.0 (WaveMetrics).

**Histology, Imaging, and Cell Counting**

Mice were deeply anesthetized with isoflurane and transcardially perfused with 50 mL of 4°C phosphate-buffered saline followed by 50 mL of 4% paraformaldehyde.

Brains were removed, post-fixed in 4% paraformaldehyde overnight at 4°C, and transferred to 30% sucrose in phosphate-buffered saline at 4°C for 3 days until they sank. Coronal sections were cut at 30 μm on a cryostat (Cryotome FSE; Thermo Fisher Scientific) and stored in antifreeze buffer at −20°C until use. Sections were counterstained with 4′,6-diamidino-2-phenylindole, mounted with DAKO mounting medium, and imaged on a confocal laser scanning microscope (LSM800; Carl Zeiss) using acquisition settings that were identical across groups. Viral expression and cannula and fiber placements were verified in every animal.

For the dual-color anterograde tracing experiment, mCherry-positive, EYFP-positive, and double-positive AHN neurons were counted within a 100 × 100 μm region of interest placed in the AHN on each of four coronal sections per animal, corresponding to anteroposterior −0.58, −0.70, −0.82, and −0.94 mm, and the four values were averaged to give a single value per animal. Counting was performed manually. A cell was scored as positive when somatic fluorescence in the corresponding channel was clearly distinguishable from the surrounding background and the labeled profile had the size and shape of a neuronal soma. A cell was scored as double labeled only when signal was present at the same position in both single-channel images and the merged image showed the combined color at that position. Images were randomized and counted by an experimenter blinded to the identity of the animal from which each image came. Because proportions are reported rather than absolute cell numbers, counting biases arising from section thickness are expected to be canceled out.

**RNA In Situ Hybridization and Atlas Registration**

A separate set of brains was processed for in situ hybridization. After perfusion, brains were post-fixed in 4% paraformaldehyde for 24 hours at 4°C and cryoprotected in 30% sucrose for 2–3 days at 4°C. Coronal sections were cut at 16 μm at −20°C on a cryostat (CM3050S; Leica Microsystems), mounted on Superfrost Plus slides (Fisherbrand), and stored overnight at −80°C.

For fluorescence imaging, slides were brought to room temperature, a drop of phosphate-buffered saline was applied to each section, and a glass coverslip was placed on top. Bright-field and mCherry fluorescence images were acquired on an SP8 confocal microscope (Leica) with a 10× objective.

In situ hybridization was performed with the RNAscope 2.5 HD Brown Chromogenic Reagent Kit (Advanced Cell Diagnostics, Cat# 322310) according to the manufacturer’s protocol, using probes against mouse Slc17a6 (Vglut2; Cat# 319171) and Slc32a1 (Vgat; Cat# 319191). Hybridization signal was developed with a 3,3′-diaminobenzidine (DAB)-based brown chromogen, and sections were counterstained with hematoxylin. Bright-field images of RNAscope-processed sections were acquired on an Axioscan 7 slide scanner (Carl Zeiss) with a 10× objective.

Fluorescence images acquired on the SP8 were registered to a reference atlas with the BigWarp plugin in ImageJ on sections corresponding to anteroposterior −0.70, −0.82, and −0.94 mm, using the anatomical boundaries of the AHN as landmarks. The position and morphological boundaries of the AHN were defined according to The Mouse Brain in Stereotaxic Coordinates (Paxinos and Franklin, 2001).

To determine the proportion of mCherry-positive AHN neurons expressing Vglut2 or Vgat, the SP8 fluorescence images and the Axioscan 7 RNAscope bright-field images were spatially aligned with BigWarp. In sections from three mice, three sections per animal, mCherry-positive cells within the AHN were counted on the aligned images, and the fraction of those cells carrying the corresponding DAB signal was calculated. As a reference population, the prevalence of the same DAB signal among all hematoxylin-stained cells within the AHN region of interest was determined in the same sections, which gives the background proportion of Vglut2- or Vgat-expressing neurons in the AHN irrespective of subicular input.

**Statistics**

Two-group comparisons used unpaired or paired two-tailed Student’s t tests, and Welch’s correction was applied when an F test indicated that the variances differed. For three or more groups, homogeneity of variance was assessed with the Brown-Forsythe test, and comparisons used ordinary one-way analysis of variance (ANOVA) with Tukey’s post hoc test when variances were homogeneous, and Brown-Forsythe and Welch ANOVA with Dunnett’s T3 post hoc test when they were not, because Tukey’s test assumes a common variance whereas Dunnett’s T3 does not. Experiments in which the same animal was tested in the absence and presence of a social target, or across successive light-stimulation epochs, were analyzed by two-way repeated-measures ANOVA followed by Šídák’s or Tukey’s multiple comparisons test. For slice electrophysiology, n denotes the number of cells, with the number of animals given alongside. The Geisser-Greenhouse correction was applied to repeated-measures analyses when sphericity was not assumed. The relationship between interaction time and distance moved was examined by linear regression, comparing slopes and then elevations between groups. For the in situ hybridization experiment, the proportion of Vglut2- or Vgat-positive cells among avSUB-recipient neurons was compared with the corresponding background prevalence in the AHN by paired two-tailed t test, with each animal contributing one pair of values. The proportions of non-GABAergic and GABAergic cells among avSUB-recipient AHN neurons were compared by paired two-tailed t test.

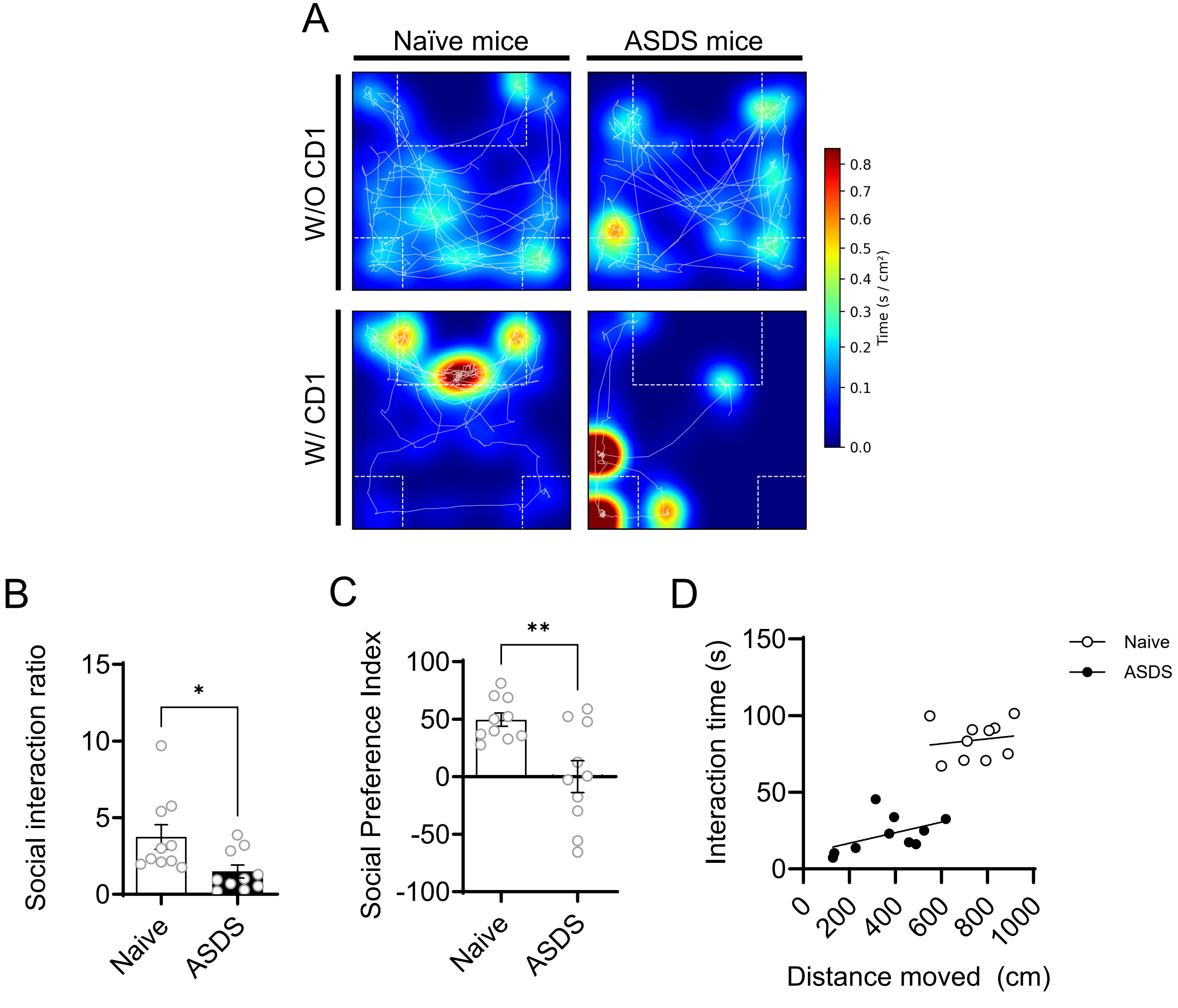
A
Naïve mice
ASDS mice
W/O CD1
W/ CD1
Time (s / cm²)
0.8
0.7
0.6
0.5
0.4
0.3
0.2
0.1
0.0
B
Social interaction ratio
15
10
5
0
*
Naive
ASDS
C
Social Preference Index
100
50
0
-50
-100
**
Naive
ASDS
D
Interaction time (s)
150
100
50
0
Distance moved (cm)
0
200
400
600
800
1000
Naive
ASDS

**Supplementary Figure S1. ASDS reduces social interaction independently of locomotion.** (A) Representative movement tracks and occupancy heat maps of a naive and a defeated mouse, without and with the CD-1 target. (B) Social interaction ratio ($p$ = .022, $n$ = 10 and 10 mice). (C) Social preference index ($p$ = .006, $n$ = 10 and 10 mice). (D) Time spent in the interaction zone with the target present plotted against distance moved. Neither slope differed from zero and the two slopes did not differ from each other, whereas the elevations differed (slopes $p$ = .669, elevations $p$ < .0001, $n$ = 10 and 10 mice). Data are mean ± SEM. Panel B was analyzed by unpaired $t$ test, panel C by unpaired $t$ test with Welch’s correction, and panel D by linear regression with comparison of slopes and elevations. Statistics are reported in Table S1.

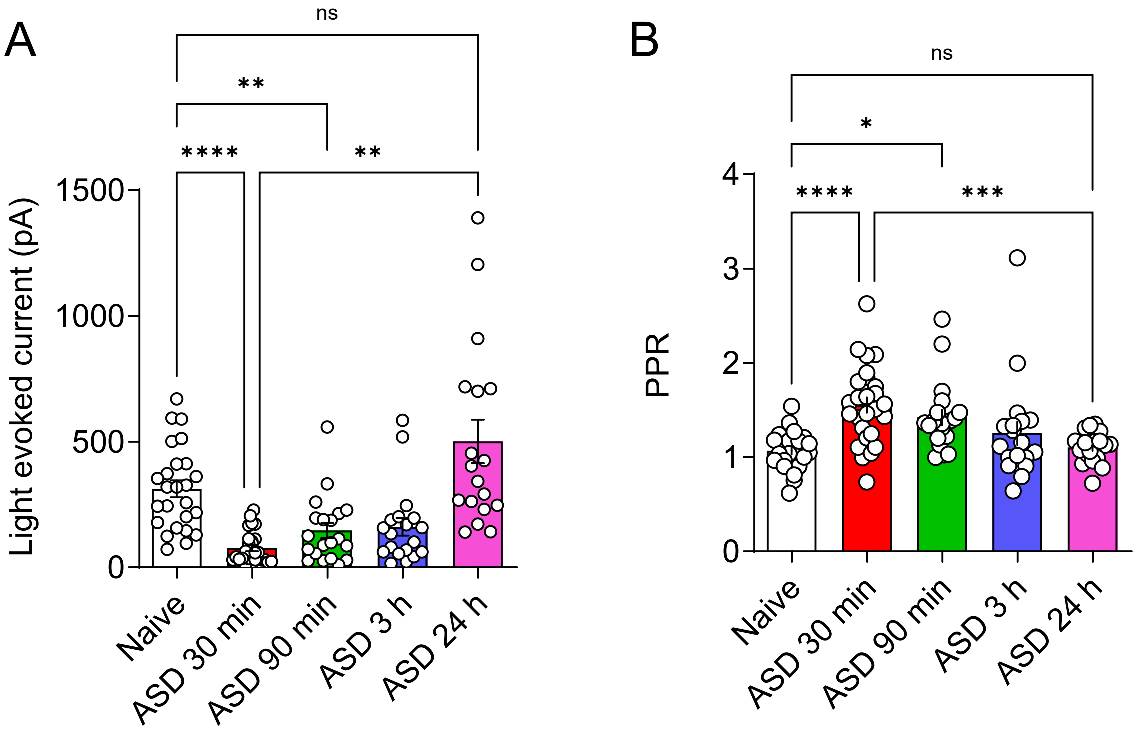

A
ns
**
****
**
1500
1000
500
0
Light evoked current (pA)
Naive
ASD 30 min
ASD 90 min
ASD 3 h
ASD 24 h
B
ns
*
****
***
4
3
2
1
0
PPR
Naive
ASD 30 min
ASD 90 min
ASD 3 h
ASD 24 h

**Supplementary Figure S2. The synaptic depression at the avSUB→AHN synapses is transient.** (A) Light-evoked current amplitude in slices prepared 30 min, 90 min, 3 hours and 24 hours after ASDS (naive versus ASDS 30 min $p < .0001$, naive versus ASDS 90 min $p < .01$, naive versus ASDS 24 hours not significant, $n = 25, 26, 20, 19$ and 18 cells from 3 mice per group). (B) Paired-pulse ratio in the same slices (naive versus ASDS 30 min $p < .0001$, naive versus ASDS 90 min $p < .05$, naive versus ASDS 24 hours not significant, $n$ as in A). Data are mean ± SEM. Panel A was analyzed by Brown-Forsythe and Welch ANOVA with Dunnett's T3 multiple comparisons test, and panel B by one-way ANOVA with Tukey's multiple comparisons test. Statistics are reported in Table S1.

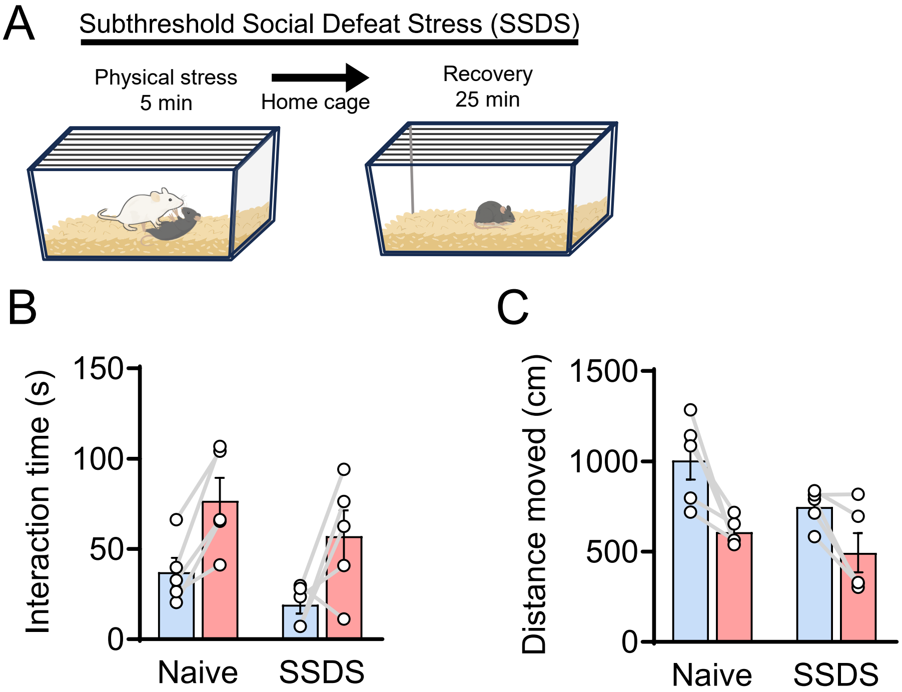
A
Subthreshold Social Defeat Stress (SSDS)
Physical stress
5 min
Home cage
Recovery
25 min
B
Interaction time (s)
150
100
50
0
Naive
SSDS
C
Distance moved (cm)
1500
1000
500
0
Naive
SSDS

**Supplementary Figure S3. SSDS does not by itself reduce social interaction.** (A) Schematic of the SSDS protocol, consisting of 5 min of physical interaction with an aggressor followed by 25 min of recovery in the home cage. (B) Time spent in the interaction zone (naive versus SSDS not significant without or with the target, $n$ = 5 and 5 mice). (C) Distance moved (naive versus SSDS not significant without or with the target, $n$ = 5 and 5 mice). Data are mean ± SEM. Both panels were analyzed by two-way repeated measures ANOVA with Šídák's multiple comparisons test. Statistics are reported in Table S1.

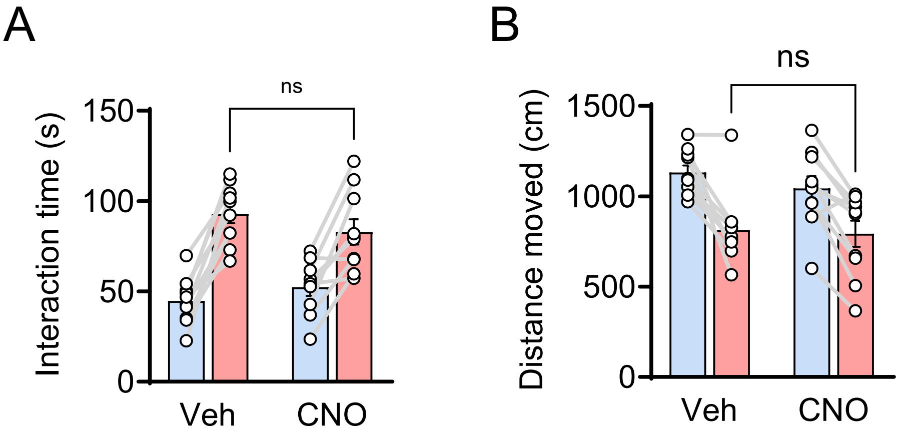
A
B
ns
ns
Interaction time (s)
150
100
50
0
Veh
CNO
Distance moved (cm)
1500
1000
500
0
Veh
CNO

**Supplementary Figure S4. Clozapine-N-oxide alone does not alter social interaction in wild-type mice.** (A) Time spent in the interaction zone in wild-type mice given vehicle or CNO (vehicle versus CNO not significant without, $p = .320$, or with, $p = .192$, the target, and both groups increased interaction time when the target was introduced, $p < .0001$ in each group, $n$ = 10 and 10 mice). (B) Distance moved in the same mice (vehicle versus CNO not significant without, $p = .319$, or with, $p = .825$, the target, $n = 10$ and 10 mice). Data are mean ± SEM. Both panels were analyzed by two-way repeated measures ANOVA with Šídák's multiple comparisons test. Statistics are reported in Table S1.

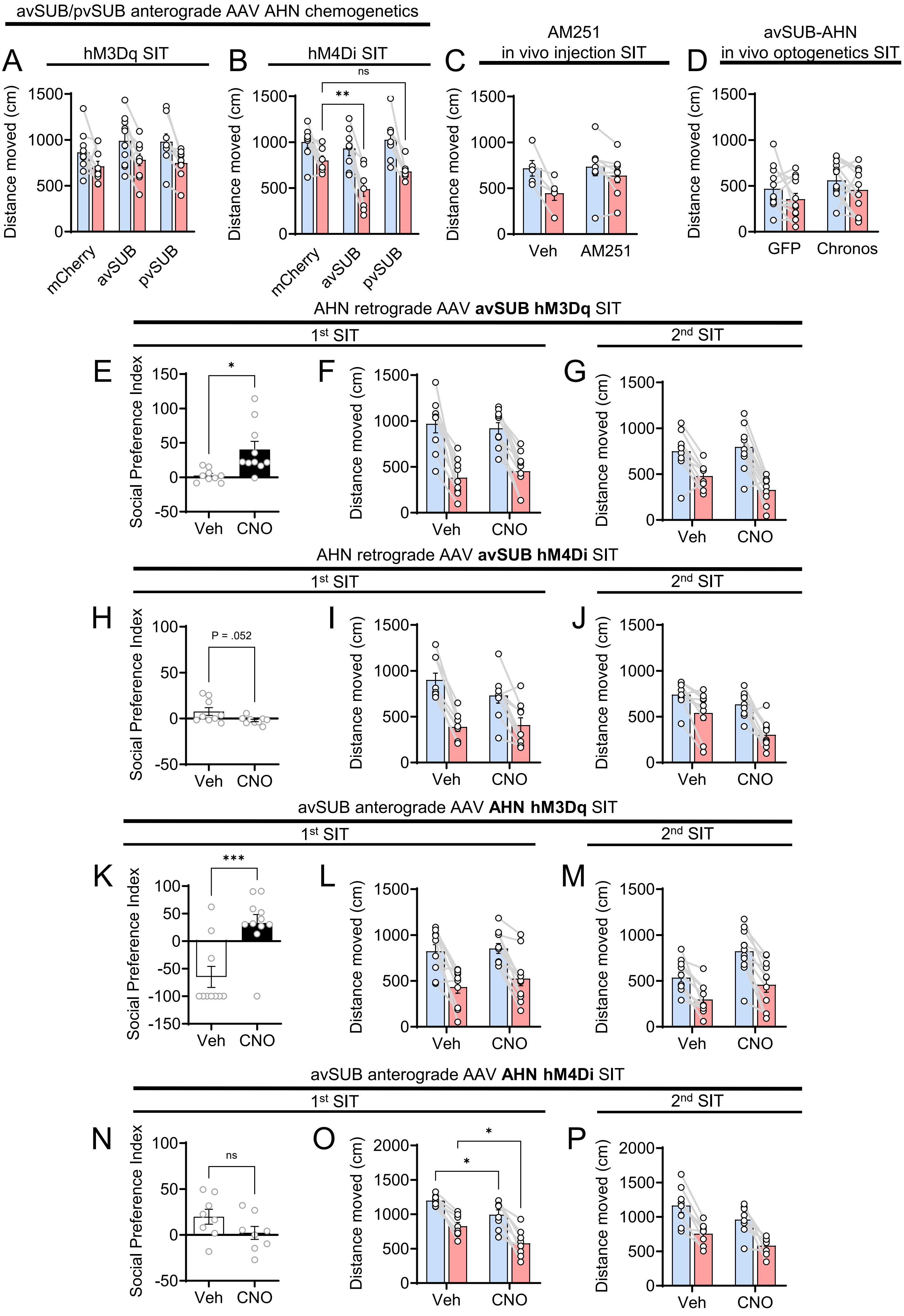
avSUB/pvSUB anterograde AAV AHN chemogenetics
A
hM3Dq SIT
B
hM4Di SIT
C
AM251
in vivo injection SIT
D
avSUB-AHN
in vivo optogenetics SIT
Distance moved (cm)
mCherry
avSUB
pvSUB
ns
**
Veh
AM251
GFP
Chronos
AHN retrograde AAV avSUB hM3Dq SIT
1st SIT
2nd SIT
E
F
G
Social Preference Index
*
CNO
AHN retrograde AAV avSUB hM4Di SIT
H
I
J
P = .052
avSUB anterograde AAV AHN hM3Dq SIT
K
L
M
***
avSUB anterograde AAV AHN hM4Di SIT
N
O
P

**Supplementary Figure S5. Social preference indices and locomotion for the chemogenetic, pharmacological and optogenetic experiments.** (A) Distance moved in stress-naive mice after activation with hM3D(Gq) (no group differences, *n* = 10 mCherry, 10 avSUB and 9 pvSUB mice). (B) Distance moved in stress-naive mice after inhibition with hM4D(Gi) (with the target present, mCherry versus avSUB *p* = .005, mCherry versus pvSUB *p* = .410, and avSUB versus pvSUB *p* = .116, and all comparisons without the target were not significant, *n* = 8 mice per group). (C) Distance moved after vehicle or AM251 infusion before ASDS (no group differences, *n* = 5 and 8 mice). (D) Distance moved in the test performed 24 hours after ASDS in the optogenetic experiment (no group differences, *n* = 12 and 12 mice). (E) Social preference index in the first test after ASDS with hM3D(Gq) in AHN-projecting avSUB neurons (vehicle versus CNO *p* = .010, *n* = 9 and 10 mice). (F) Distance moved in the first test (no group differences, *n* = 9 and 10 mice). (G) Distance moved in the second test 24 hours later (no group differences, *n* as in F). (H) Social preference index in the first test after SSDS with hM4D(Gi) in the same population (vehicle versus CNO *p* = .052, *n* = 9 and 9 mice). (I) Distance moved in the first test (no group differences, *n* = 9 and 9 mice). (J) Distance moved in the second test 24 hours later (no group differences, *n* as in I). (K) Social preference index in the first test after ASDS with hM3D(Gq) in AHN neurons receiving avSUB input (vehicle versus CNO *p* < .001, *n* = 10 and 11 mice). (L) Distance moved in the first test (no group differences, *n* = 10 and 11 mice). (M) Distance moved in the second test 24 hours later (no group differences, *n* as in L). (N) Social preference index in the first test after SSDS with hM4D(Gi) in the same population (vehicle versus CNO *p* = .123, *n* = 8 and 8 mice). (O) Distance moved in the first test (vehicle versus CNO *p* = .038 without and *p* = .011 with the target, *n* = 8 and 8 mice). (P) Distance moved in the second test 24 hours later (no group differences, *n* = 8 and 7 mice). Data are mean ± SEM. The distance-moved panels were analyzed by two-way repeated measures ANOVA with Šídák’s or Tukey’s multiple comparisons test, and the social preference index panels by unpaired *t* test with Welch’s correction, except panel N, which used an unpaired *t* test. Statistics are reported in Table S1.

**Supplemental Table S1. Statistical analyses**

Two-tailed tests throughout, significance set at p < .05. For slice electrophysiology, n is given as cells (animals).

| Figure | Panel | Measure | Comparison | Test | Statistic | p | Summary | n |
|---|---|---|---|---|---|---|---|---|
| Figure 1 | b | Interaction time | Stress x target (interaction) | Two-way RM ANOVA | F(1,18) = 46.19 | <.0001 | **** | Naive, 10 mice<br>ASDS, 10 mice |
| | | | Stress (naive vs ASDS) | | F(1,18) = 52.56 | <.0001 | **** | |
| | | | Target (W/O vs W/) | | F(1,18) = 40.42 | <.0001 | **** | |
| | | | Naive vs ASDS, W/O | Šídák's | t = 0.93 | .587 | ns | |
| | | | Naive vs ASDS, W/ | Šídák's | t = 9.93 | <.0001 | **** | |
| Figure 1 | c | Corner duration | Stress x target (interaction) | Two-way RM ANOVA | F(1,18) = 39.74 | <.0001 | **** | Naive, 10 mice<br>ASDS, 10 mice |
| | | | Stress (naive vs ASDS) | | F(1,18) = 41.07 | <.0001 | **** | |
| | | | Target (W/O vs W/) | | F(1,18) = 1.85 | .191 | ns | |
| | | | Naive vs ASDS, W/O | Šídák's | t = 1.59 | .225 | ns | |
| | | | Naive vs ASDS, W/ | Šídák's | t = 8.87 | <.0001 | **** | |
| Figure 1 | d | Distance moved | Stress x target (interaction) | Two-way RM ANOVA | F(1,18) = 5.96 | .025 | * | Naive, 10 mice<br>ASDS, 10 mice |
| | | | Stress (naive vs ASDS) | | F(1,18) = 18.10 | .0005 | *** | |
| | | | Target (W/O vs W/) | | F(1,18) = 81.89 | <.0001 | **** | |
| | | | Naive vs ASDS, W/O | Šídák's | t = 2.23 | .063 | ns | |
| | | | Naive vs ASDS, W/ | Šídák's | t = 4.90 | <.0001 | **** | |
| Figure 1 | e | Interaction time, 24 h after ASDS | Stress x target (interaction) | Two-way RM ANOVA | F(1,18) = 5.89 | .026 | * | Naive, 10 mice<br>ASDS, 10 mice |
| | | | Stress (naive vs ASDS) | | F(1,18) = 20.34 | .0003 | *** | |
| | | | Target (W/O vs W/) | | F(1,18) = 21.34 | .0002 | *** | |
| | | | Naive vs ASDS, W/O | Šídák's | t = 2.02 | .099 | ns | |
| | | | Naive vs ASDS, W/ | Šídák's | t = 5.04 | <.0001 | **** | |
| Figure 1 | h | Interaction time (hM3Dq) | Group x target (interaction) | Two-way RM ANOVA | F(2,26) = 0.62 | .544 | ns | mCherry, 10 mice<br>avSUB, 10 mice<br>pvSUB, 9 mice |
| | | | Group | | F(2,26) = 2.36 | .115 | ns | |
| | | | Target | | F(1,26) = 180.4 | <.0001 | **** | |
| | | | mCherry vs avSUB, W/ | Tukey's | q = 1.95 | .359 | ns | |
| | | | mCherry vs pvSUB, W/ | Tukey's | q = 3.07 | .086 | ns | |
| | | | avSUB vs pvSUB, W/ | Tukey's | q = 1.17 | .688 | ns | |
| Figure 1 | i | Interaction time (hM4Di) | Group x target (interaction) | Two-way RM ANOVA | F(2,21) = 7.07 | .005 | ** | mCherry, 8 mice<br>avSUB, 8 mice<br>pvSUB, 8 mice |
| | | | Group | | F(2,21) = 5.99 | .009 | ** | |
| | | | Target | | F(1,21) = 63.87 | <.0001 | **** | |
| | | | mCherry vs avSUB, W/ | Tukey's | q = 5.19 | .002 | ** | |
| | | | mCherry vs pvSUB, W/ | Tukey's | q = 0.68 | .881 | ns | |
| | | | avSUB vs pvSUB, W/ | Tukey's | q = 5.87 | .0005 | *** | |
| Figure 2 | b | Light-evoked EPSC (avSUB) | Among groups | Brown-Forsythe ANOVA | F*(2, 31.26) = 13.03 | <.0001 | **** | Naive, 12 cells from 5 mice<br>ASDS, 26 cells from 7 mice<br>RU486+ASDS, 21 cells from 4 mice |
| | | | Among groups | Welch's ANOVA | W(2, 20.91) = 18.79 | <.0001 | **** | |
| | | | Naive vs ASDS | Dunnett's T3 | t = 4.56 | .002 | ** | |
| | | | Naive vs RU486+ASDS | Dunnett's T3 | t = 0.45 | .958 | ns | |
| | | | ASDS vs RU486+ASDS | Dunnett's T3 | t = 4.48 | .0005 | *** | |
| Figure 2 | c | Paired-pulse ratio (avSUB) | Among groups | Brown-Forsythe ANOVA | F*(2, 53.80) = 23.97 | <.0001 | **** | Naive, 12 cells from 5 mice<br>ASDS, 26 cells from 7 mice<br>RU486+ASDS, 21 cells from 4 mice |
| | | | Among groups | Welch's ANOVA | W(2, 34.31) = 17.02 | <.0001 | **** | |
| | | | Naive vs ASDS | Dunnett's T3 | t = 4.39 | .0003 | *** | |
| | | | Naive vs RU486+ASDS | Dunnett's T3 | t = 1.60 | .316 | ns | |
| | | | ASDS vs RU486+ASDS | Dunnett's T3 | t = 5.83 | <.0001 | **** | |

**Supplemental Table S1. Statistical analyses**

Two-tailed tests throughout, significance set at p < .05. For slice electrophysiology, n is given as cells (animals).

| Figure | Panel | Measure | Comparison | Test | Statistic | p | Summary | n |
|---|---|---|---|---|---|---|---|---|
| Figure 2 | e | Light-evoked EPSC (pvSUB) | Naive vs ASDS | Unpaired t test, Welch's correction | t(23.28) = 1.49 | .150 | ns | Naive, 17 cells from 3 mice<br>ASDS, 17 cells from 4 mice |
| Figure 2 | f | Paired-pulse ratio (pvSUB) | Naive vs ASDS | Unpaired t test, Welch's correction | t(31.18) = 0.39 | .699 | ns | Naive, 17 cells from 3 mice<br>ASDS, 17 cells from 4 mice |
| Figure 3 | a | Light-evoked EPSC | Vehicle vs CORT | Brown-Forsythe and Welch ANOVA, Dunnett's T3 | t = 4.46 | .001 | ** | Vehicle, 13 cells from 5 mice<br>Corticosterone, 13 cells from 4 mice<br>AM251+corticosterone, 15 cells from 4 mice |
| | | | Vehicle vs AM251+CORT | Dunnett's T3 | t = 0.89 | .754 | ns | |
| | | | CORT vs AM251+CORT | Dunnett's T3 | t = 3.90 | .003 | ** | |
| Figure 3 | b | Paired-pulse ratio | Vehicle vs CORT | Brown-Forsythe and Welch ANOVA, Dunnett's T3 | t = 3.19 | .019 | * | Vehicle, 13 cells from 5 mice<br>Corticosterone, 13 cells from 4 mice<br>AM251+corticosterone, 15 cells from 4 mice |
| | | | Vehicle vs AM251+CORT | Dunnett's T3 | t = 0.16 | .998 | ns | |
| | | | CORT vs AM251+CORT | Dunnett's T3 | t = 2.95 | .023 | * | |
| Figure 3 | c | Light-evoked EPSC | Baseline vs ACEA 30 min | Paired t test | t(6) = 2.88 | .028 | * | 7 cells from 4 mice, each recorded before and after ACEA |
| Figure 3 | d | Paired-pulse ratio | Baseline vs ACEA 30 min | Paired t test | t(6) = 3.39 | .015 | * | 7 cells from 4 mice, each recorded before and after ACEA |
| Figure 3 | f | Interaction time | Treatment x target (interaction) | Two-way RM ANOVA | F(1,11) = 7.07 | .022 | * | Vehicle, 5 mice<br>AM251, 8 mice |
| | | | Vehicle vs AM251, W/O | Šídák's | t = 0.04 | .999 | ns | |
| | | | Vehicle vs AM251, W/ | Šídák's | t = 3.25 | .007 | ** | |
| Figure 3 | g | Light-evoked EPSC | Vehicle vs AM251 | Unpaired t test, Welch's correction | t(30.80) = 2.99 | .006 | ** | Vehicle, 27 cells from 4 mice<br>AM251, 25 cells from 4 mice |
| Figure 3 | h | Paired-pulse ratio | Vehicle vs AM251 | Unpaired t test, Welch's correction | t(37.70) = 3.22 | .003 | ** | Vehicle, 27 cells from 4 mice<br>AM251, 25 cells from 4 mice |
| Figure 4 | b | Interaction time, 1st SIT | Vehicle vs CNO, W/ | Two-way RM ANOVA, Šídák's | t = 4.45 | .0002 | *** | Vehicle, 9 mice<br>CNO, 10 mice |
| Figure 4 | c | Interaction time, 2nd SIT | Vehicle vs CNO, W/ | Two-way RM ANOVA, Šídák's | t = 2.97 | .011 | * | Vehicle, 9 mice<br>CNO, 10 mice |
| Figure 4 | d | Interaction time, 1st SIT | Vehicle vs CNO, W/ | Two-way RM ANOVA, Šídák's | t = 2.96 | .012 | * | Vehicle, 9 mice<br>CNO, 9 mice |
| Figure 4 | e | Interaction time, 2nd SIT | Vehicle vs CNO, W/ | Two-way RM ANOVA, Šídák's | t = 3.19 | .006 | ** | Vehicle, 9 mice<br>CNO, 9 mice |
| Figure 4 | g | Interaction time, 1st SIT | Vehicle vs CNO, W/ | Two-way RM ANOVA, Šídák's | t = 3.84 | .0009 | *** | Vehicle, 10 mice<br>CNO, 11 mice |
| Figure 4 | h | Interaction time, 2nd SIT | Vehicle vs CNO, W/ | Two-way RM ANOVA, Šídák's | t = 4.30 | .0002 | *** | Vehicle, 10 mice<br>CNO, 11 mice |
| Figure 4 | i | Interaction time, 1st SIT | Vehicle vs CNO, W/ | Two-way RM ANOVA, Šídák's | t = 4.83 | <.0001 | **** | Vehicle, 8 mice<br>CNO, 8 mice |
| | | | Vehicle vs CNO, W/O | Šídák's | t = 2.65 | .026 | * | |
| Figure 4 | j | Interaction time, 2nd SIT | Vehicle vs CNO, W/ | Two-way RM ANOVA, Šídák's | t = 2.78 | .020 | * | Vehicle, 8 mice<br>CNO, 7 mice |
| Figure 5 | c | Interaction time across epochs | Group x epoch (interaction) | Two-way RM ANOVA | F(4,84) = 2.45 | .052 | ns | GFP, 11 mice<br>Chronos, 12 mice |
| | | | Epoch | | F(3.145, 66.05) = 5.22 | .002 | ** | |
| | | | Group (GFP vs Chronos) | | F(1,21) = 8.46 | .008 | ** | |
| | | | GFP vs Chronos, 1st light-on | Šídák's | t = 3.68 | .010 | * | |
| | | | GFP vs Chronos, 2nd light-on | Šídák's | t = 2.83 | .052 | ns | |
| Figure 5 | d | Interaction time across epochs | Group x epoch (interaction) | Two-way RM ANOVA | F(4,84) = 2.41 | .056 | ns | GFP, 11 mice<br>Chronos, 12 mice |
| | | | Group (GFP vs Chronos) | | F(1,21) = 8.97 | .007 | ** | |
| | | | GFP vs Chronos, 1st light-on | Šídák's | t = 3.37 | .016 | * | |
| | | | GFP vs Chronos, 2nd light-on | Šídák's | t = 3.09 | .038 | * | |
| Figure 5 | e | Interaction time, 24 h after ASDS | GFP vs Chronos, W/O | Two-way RM ANOVA, Šídák's | t = 1.82 | .147 | ns | GFP, 12 mice<br>Chronos, 12 mice |

**Supplemental Table S1. Statistical analyses**

Two-tailed tests throughout, significance set at p < .05. For slice electrophysiology, n is given as cells (animals).

| Figure | Panel | Measure | Comparison | Test | Statistic | p | Summary | n |
|---|---|---|---|---|---|---|---|---|
| | | | GFP vs Chronos, W/ | Šídák's | t = 0.78 | .688 | ns | |
| Figure 6 | g | Non-GABAergic vs GABAergic among avSUB-recipient AHN neurons | GABAergic vs non-GABAergic | Paired t test | t(3) = 5.45 | .012 | * | 4 mice, each contributing one pair of values |
| Figure 6 | i | VGLUT2-positive cells (%) | avSUB-recipient vs all AHN neurons | Paired t test | t(2) = 5.31 | .034 | * | 3 mice, each contributing one pair of values |
| Figure 6 | j | VGAT-positive cells (%) | avSUB-recipient vs all AHN neurons | Paired t test | t(2) = 2.13 | .167 | ns | 3 mice, each contributing one pair of values |
| Figure 7 | b | OFT center duration (hM3Dq), W/O ASDS | Omnibus | Welch's ANOVA | W(2, 13.83) = 12.13 | 0.0009 | *** | mCherry, 9 mice<br>avSUB, 9 mice<br>pvSUB, 10 mice |
| | | | mCherry vs avSUB | Dunnett's T3 | t = 1.036 | 0.6666 | ns | |
| | | | mCherry vs pvSUB | Dunnett's T3 | t = 3.435 | 0.0211 | * | |
| | | | avSUB vs pvSUB | Dunnett's T3 | t = 4.144 | 0.0034 | ** | |
| Figure 7 | c | OFT center duration (hM4Di), W/O ASDS | Omnibus | One-way ANOVA | F(2,21) = 3.294 | 0.057 | ns | mCherry, 8 mice<br>avSUB, 8 mice<br>pvSUB, 8 mice |
| Figure 7 | d | EPM open arm duration (hM3Dq), W/O ASDS | Omnibus | One-way ANOVA | F(2,27) = 11.41 | 0.0003 | *** | mCherry, 10 mice<br>avSUB, 10 mice<br>pvSUB, 10 mice |
| | | | mCherry vs avSUB | Tukey's | q = 3.12 | 0.0884 | ns | |
| | | | mCherry vs pvSUB | Tukey's | q = 3.63 | 0.0413 | * | |
| | | | avSUB vs pvSUB | Tukey's | q = 6.75 | 0.0002 | *** | |
| Figure 7 | e | EPM closed arm duration (hM3Dq), W/O ASDS | Omnibus | One-way ANOVA | F(2,27) = 18.30 | <.0001 | **** | mCherry, 10 mice<br>avSUB, 10 mice<br>pvSUB, 10 mice |
| | | | mCherry vs avSUB | Tukey's | q = 2.89 | 0.1211 | ns | |
| | | | mCherry vs pvSUB | Tukey's | q = 5.53 | 0.0016 | ** | |
| | | | avSUB vs pvSUB | Tukey's | q = 8.42 | <.0001 | **** | |
| Figure 7 | f | EPM open arm duration (hM4Di), W/O ASDS | Omnibus | One-way ANOVA | F(2,21) = 4.982 | 0.017 | * | mCherry, 8 mice<br>avSUB, 8 mice<br>pvSUB, 8 mice |
| | | | mCherry vs avSUB | Tukey's | q = 0.62 | 0.9001 | ns | |
| | | | mCherry vs pvSUB | Tukey's | q = 4.14 | 0.0211 | * | |
| | | | avSUB vs pvSUB | Tukey's | q = 3.52 | 0.0534 | ns | |
| Figure 7 | g | EPM closed arm duration (hM4Di), W/O ASDS | Omnibus | One-way ANOVA | F(2,21) = 4.645 | 0.0214 | * | mCherry, 8 mice<br>avSUB, 8 mice<br>pvSUB, 8 mice |
| | | | mCherry vs avSUB | Tukey's | q = 0.08 | 0.9982 | ns | |
| | | | mCherry vs pvSUB | Tukey's | q = 3.77 | 0.0368 | * | |
| | | | avSUB vs pvSUB | Tukey's | q = 3.69 | 0.0415 | * | |
| Figure 7 | h | Social preference index (hM3Dq) | Omnibus | Welch's ANOVA | W(2, 13.24) = 22.41 | <.0001 | **** | mCherry, 10 mice<br>avSUB, 10 mice<br>pvSUB, 9 mice |
| | | | mCherry vs avSUB | Dunnett's T3 | t = 5.304 | 0.0006 | *** | |
| | | | mCherry vs pvSUB | Dunnett's T3 | t = 0.884 | 0.7606 | ns | |
| | | | avSUB vs pvSUB | Dunnett's T3 | t = 4.803 | 0.0028 | ** | |
| Figure 7 | i | Interaction time (hM3Dq) | mCherry vs avSUB, W/ | Two-way RM ANOVA, Tukey's | t = 3.960 | 0.0007 | *** | mCherry, 10 mice<br>avSUB, 10 mice<br>pvSUB, 9 mice |
| | | | avSUB vs pvSUB, W/ | Tukey's | t = 4.044 | 0.0005 | *** | |
| | | | mCherry vs pvSUB, W/ | Tukey's | t = 0.190 | 0.9966 | ns | |
| Figure 7 | j | Social preference index (hM4Di) | Omnibus | Brown-Forsythe ANOVA | F*(2, 16.54) = 0.978 | 0.3968 | ns | mCherry, 8 mice<br>avSUB, 8 mice<br>pvSUB, 8 mice |
| | | | mCherry vs avSUB | Dunnett's T3 | t = 1.217 | 0.5511 | ns | |
| | | | mCherry vs pvSUB | Dunnett's T3 | t = 0.610 | 0.9018 | ns | |
| | | | avSUB vs pvSUB | Dunnett's T3 | t = 0.951 | 0.7202 | ns | |
| Figure 7 | k | Interaction time (hM4Di) | All comparisons, W/ | Two-way RM ANOVA, Tukey's | — | ≥.5046 | ns | mCherry, 8 mice<br>avSUB, 8 mice<br>pvSUB, 8 mice |
| Figure S1 | b | Social interaction ratio | Naive vs ASDS | Unpaired t test | t(18) = 2.50 | .022 | * | Naive, 10 mice<br>ASDS, 10 mice |
| Figure S1 | c | Social preference index | Naive vs ASDS | Unpaired t test, Welch's correction | t(12.06) = 3.30 | .006 | ** | Naive, 10 mice<br>ASDS, 10 mice |

**Supplemental Table S1. Statistical analyses**

Two-tailed tests throughout, significance set at p < .05. For slice electrophysiology, n is given as cells (animals).

| Figure | Panel | Measure | Comparison | Test | Statistic | p | Summary | n |
|---|---|---|---|---|---|---|---|---|
| Figure S1 | d | Interaction time vs distance moved | Slope, naive | Linear regression | F(1,8) = 0.19 | .677 | ns | Naive, 10 mice<br>ASDS, 10 mice |
| | | | Slope, ASDS | Linear regression | F(1,8) = 2.35 | .164 | ns | |
| | | | Slopes, naive vs ASDS | Comparison of slopes | F(1,16) = 0.19 | .669 | ns | |
| | | | Elevations, naive vs ASDS | Comparison of elevations | F(1,17) = 30.25 | <.0001 | **** | |
| Figure S2 | a | Light-evoked EPSC | Among groups | Brown-Forsythe ANOVA | F*(4, 36.66) = 13.67 | <.0001 | **** | Naive, 25 cells from 3 mice<br>ASDS 30 min, 26 cells from 3 mice<br>ASDS 90 min, 20 cells from 3 mice<br>ASDS 3 h, 19 cells from 3 mice<br>ASDS 24 h, 18 cells from 3 mice |
| | | | Among groups | Welch's ANOVA | W(4, 43.99) = 15.87 | <.0001 | **** | |
| | | | Naive vs ASDS 30 min | Dunnett's T3 | t = 6.61 | <.0001 | **** | |
| | | | Naive vs ASDS 90 min | Dunnett's T3 | t = 3.73 | <.01 | ** | |
| | | | Naive vs ASDS 24 h | Dunnett's T3 | t = 2.03 | ns | ns | |
| | | | ASDS 30 min vs ASDS 24 h | Dunnett's T3 | t = 4.85 | <.01 | ** | |
| Figure S2 | b | Paired-pulse ratio | Among groups | One-way ANOVA | F(4,103) = 7.61 | <.0001 | **** | Naive, 25 cells from 3 mice<br>ASDS 30 min, 26 cells from 3 mice<br>ASDS 90 min, 20 cells from 3 mice<br>ASDS 3 h, 19 cells from 3 mice<br>ASDS 24 h, 18 cells from 3 mice |
| | | | Naive vs ASDS 30 min | Tukey's | q = 6.73 | <.0001 | **** | |
| | | | Naive vs ASDS 90 min | Tukey's | q = 4.58 | <.05 | * | |
| | | | Naive vs ASDS 24 h | Tukey's | q = 0.44 | ns | ns | |
| | | | ASDS 30 min vs ASDS 24 h | Tukey's | q = 5.70 | <.001 | *** | |
| Figure S3 | b | Interaction time | Naive vs SSDS, W/O | Two-way RM ANOVA, Šídák's | t = 1.82 | .165 | ns | Naive, 5 mice<br>SSDS, 5 mice |
| | | | Naive vs SSDS, W/ | Šídák's | t = 1.82 | .165 | ns | |
| Figure S3 | c | Distance moved | Naive vs SSDS, W/O | Two-way RM ANOVA, Šídák's | t = 2.29 | .069 | ns | Naive, 5 mice<br>SSDS, 5 mice |
| | | | Naive vs SSDS, W/ | Šídák's | t = 2.29 | .069 | ns | |
| Figure S4 | a | Interaction time | Vehicle vs CNO, W/O | Two-way RM ANOVA, Šídák's | t = 1.01 | .320 | ns | Vehicle, 10 mice<br>CNO, 10 mice |
| | | | Vehicle vs CNO, W/ | Šídák's | t = 1.33 | .192 | ns | |
| | | | W/O vs W/, vehicle | Šídák's | t = 8.26 | <.0001 | **** | |
| | | | W/O vs W/, CNO | Šídák's | t = 5.25 | <.0001 | **** | |
| Figure S4 | b | Distance moved | Vehicle vs CNO, W/O | Two-way RM ANOVA, Šídák's | t = 1.01 | .319 | ns | Vehicle, 10 mice<br>CNO, 10 mice |
| | | | Vehicle vs CNO, W/ | Šídák's | t = 0.22 | .825 | ns | |
| Figure S5 | b | Distance moved (hM4Di) | mCherry vs avSUB, W/ | Two-way RM ANOVA, Tukey's | q = 4.70 | .005 | ** | mCherry, 8 mice<br>avSUB, 8 mice<br>pvSUB, 8 mice |
| | | | mCherry vs pvSUB, W/ | Tukey's | q = 1.82 | .410 | ns | |
| | | | avSUB vs pvSUB, W/ | Tukey's | q = 2.88 | .116 | ns | |
| Figure S5 | e | Social preference index | Vehicle vs CNO | Unpaired t test, Welch's correction | t(10.25) = 3.15 | .010 | * | Vehicle, 9 mice<br>CNO, 10 mice |
| Figure S5 | h | Social preference index | Vehicle vs CNO | Unpaired t test, Welch's correction | t(9.64) = 2.22 | .052 | ns | Vehicle, 9 mice<br>CNO, 9 mice |
| Figure S5 | k | Social preference index | Vehicle vs CNO | Unpaired t test, Welch's correction | t(17.68) = 4.00 | <.001 | *** | Vehicle, 10 mice<br>CNO, 11 mice |
| Figure S5 | n | Social preference index | Vehicle vs CNO | Unpaired t test | t(14) = 1.64 | .123 | ns | Vehicle, 8 mice<br>CNO, 8 mice |
| Figure S5 | o | Distance moved | Treatment x target (interaction) | Two-way RM ANOVA | F(1,14) = 0.25 | .627 | ns | Vehicle, 8 mice<br>CNO, 8 mice |
| | | | Treatment (vehicle vs CNO) | | F(1,14) = 10.40 | .006 | ** | |
| | | | Vehicle vs CNO, W/O | Šídák's | t = 2.49 | .038 | * | |
| | | | Vehicle vs CNO, W/ | Šídák's | t = 3.01 | .011 | * | |
